# Therapeutic CD8+T cell tissue retention and immunomodulation during ART interruption fails to prevent SIV rebound

**DOI:** 10.1101/2025.03.17.643755

**Authors:** M. Betina Pampena, Sadia Samer, Elise G. Viox, Kevin Nguyen, Claire Deleage, Leticia Kuri-Cervantes, James Regan, Vincent H. Wu, Steffen Docken, Jeffrey T. Safrit, Katharine J. Bar, Brandon F. Keele, Miles P. Davenport, Mirko Paiardini, Michael R. Betts

**Affiliations:** Department of Microbiology, Perelman School of Medicine, University of Pennsylvania, Philadelphia, PA, USA; Institute for Immunology, Perelman School of Medicine, University of Pennsylvania, Philadelphia, PA, USA; Division of Microbiology and Immunology, Emory National Primate Research Center, Emory University, Atlanta, GA, USA; AIDS and Cancer Virus Program, Frederick National Laboratory for Cancer Research, Acquired Immunodeficiency Syndrome (AIDS) and Cancer Virus Program, Leidos Biomedical Research Inc., Frederick, MD, USA; Kirby Institute for Infection and Immunity, University of New South Wales, Sydney, NSW, Australia; ImmunityBio, Inc., Culver City, CA, USA; Department of Pathology and Laboratory Medicine, School of Medicine, Emory University, Atlanta, GA 30322, USA

## Abstract

A primary obstacle for HIV elimination is the long-term viral reservoir in lymphoid tissues (LT) that can cause rebound viremia if therapy is stopped. Cytotoxic CD8+ T cells are critical for control of HIV and SIV viremia; however, CD8+ T cells that migrate to LT are primarily non-cytotoxic, calling into question whether these cells could reduce the viral reservoir on ART or control viral replication when therapy is halted. To determine if CD8+ T cells can inhibit viral replication when retained in lymphoid tissues, we inhibited lymphocyte egress from LTs in antiretroviral therapy (ART)-treated SIV-infected rhesus macaques (RM) during analytic treatment interruption (ATI) using the S1PR modulator FTY720 alone or in combination with anti-PD1 and the IL-15 receptor superagonist N-803 to increase cytolytic function. FTY720 retained migrating CD4+ and CD8+ T cells in LT, whereas cytotoxic CD8+ T cells remained in the vasculature. After ATI and viral rebound, activated SIV-specific CD8+ T cells increased in frequency in LT of FTY720-treated RM but failed to become cytotoxic or control plasma viremia compared to controls, even when combined with anti-PD1 and N-803. These findings indicate that LT-localized CD8+ T cells alone are insufficient to delay or prevent plasma viral rebound during ATI.

**Significance Statement:** HIV therapeutic and cure strategies have all operated under the assumption that CD8+ T cells can become cytotoxic in lymph nodes to eliminate infected cells from reservoir repositories. Using FTY720 to prevent lymphocyte egress from lymph nodes, we demonstrated that lymphoid tissue-restricted CD8+ T cells are insufficient to prevent SIV rebound during antiretroviral treatment interruption, even in the presence of immunomodulators such as IL-15 receptor superagonist and PD-1 blockade. These findings call into question the ability of lymph node CD8+ T cells to control or eliminate SIV/HIV infection from lymphoid tissues without substantial immunomodulation, a notion contrary to current efforts to engage CD8+ T cells for therapeutic elimination of HIV viral reservoirs during ART.

## INTRODUCTION

Elimination of the latent HIV reservoir, composed primarily of infected CD4+ T cells, remains the key challenge for immune mediated clearance of HIV infection and the focus of HIV cure-based research. While antiretroviral treatment (ART) can suppress viral replication, the long-lived nature and homeostatic renewal capability of infected CD4+ T cells perpetuates the viral reservoir for the lifetime of the host (1). The HIV-infected CD4+ T cell reservoir resides predominately in lymphoid tissues (lymph node (LN)- and gut-associated lymphoid tissues) irrespective of treatment status (2–4). Interestingly, in the absence of therapy, T follicular helper cells (TFH) serve as the dominant cell type for viral infection, replication, and viral production; and during ART, they are responsible for persistent HIV-1 transcription within lymphoid tissue (5, 6). Indeed, in the Simian Immunodeficiency Virus (SIV) rhesus macaque (RM) model of HIV infection, it was shown that >98% of SIV-infected cells are found within gut and lymphoid tissues (2, 7). However, many infected CD4+ T cells in tissues and peripheral blood have migratory and tissue homing properties, and are therefore capable of continual redistribution across the body (1). Development of strategies to clear the HIV reservoir are paramount to HIV eradication efforts.

Numerous observations have supported the key role of CD8+ T cells in the control of HIV infection and disease progression, including the temporal association between HIV-specific CD8+ T cell expansion and control of acute viremia(8, 9), HIV-specific CD8+ T cell functional properties and non-progression (10, 11), and loss of disease control after viral escape within targeted CD8+ T cell epitopes (12, 13). Non-human primate (NHP) studies have further supported the role of CD8+ T cells, showing heightened viral load in SIV-infected macaques after CD8+ T cell depletion and a particular requirement of these cells for maintaining therapy-induced viral suppression (14–16). Additionally, in SIV-infected Mamu B*08/B*17 controller RMs, control of viral replication in LN T cell zones is abrogated after CD8 depletion (17). Furthermore, we have recently shown evidence that SIV-specific CD8+ T cells in Mamu A*01 RMs can inhibit the rebound of susceptible virus following treatment interruption, thereby having a ‘sieving’ effect on the rebounding virus (18).

Based on work studying peripheral blood CD8+ T cell responses, cytotoxic activity mediated by perforin/granzyme B (8, 19–22) has been associated with HIV immune control. However, fully differentiated cytotoxic CD8+ T cell subsets are largely restricted to the vasculature (including spleen and bone marrow) due in part to the absence of CCR7 and CD62L expression(23, 24). Thus, their role in controlling lymphoid tissue HIV reservoirs is unclear. We and others have shown that cytotoxic CD8 activity is lost in LN rapidly after acute SIV/HIV infection (25, 26), and remains significantly lower than blood during chronic infection irrespective of treatment or progression status (26–29). However, despite the lack of cytotoxicity, HIV- and SIV-specific CD8+ T cells can readily be observed in lymphoid tissues (either as migrating or resident cells) during all phases of infection (25–28, 30, 31). While it remains unclear what role tissue-homing or tissue-resident CD8+ T cells play in controlling plasma viremia, it seems evident that migration of infected CD4+ T cells into lymphoid tissues facilitates the circumvention of surveillance by blood cytotoxic CD8+ T cells.

To directly address this issue, we used the lymphocyte migration inhibitor Fingolimod (FTY720) alone or in combination with immunomodulators αPD1 and the IL-15 receptor superagonist (IL15R SA) N-803 to inhibit lymphocyte egress from lymphoid tissues during analytic therapy interruption (ATI) in SIV infected RM and increase cytolytic properties of effector cells. FTY720 is a sphingosine-1 phosphate receptor (S1PR) modulator that blocks the interaction of S1P with four of its receptors (S1PR1, S1PR3, S1PR4 and S1PR5), the major pathway involved in lymphocyte migration (32). FTY720 has previously been used in ART-suppressed, SIV-infected RMs where it resulted in a decrease of SIV DNA content in blood as well as in LN follicular helper T cells in most treated animals (33). Based upon this, we hypothesized that FTY720-induced migration inhibition would trap migrating infected CD4+ T cells in secondary lymphoid tissues, allowing us to determine the differential role of vascular-restricted vs. tissue homing and resident CD8+ T cells in the control of SIV replication. We further hypothesized that SIV-specific CD8+ T cells trapped in LN would acquire cytotoxic function in response to SIV reactivation and the performed immune-based interventions, potentially leading to enhanced viral control.

While we found that FTY720 effectively redistributed migrating T cells into tissues, including both CD4+ and CD8+ T cells, the fully differentiated cytotoxic T cell subsets remained in the vasculature. Contrary to our expectations, LN SIV-specific CD8+ T cells in FTY720-treated animals, irrespective of the addition of αPD1 and N-803, did not acquire cytotoxic properties during ATI, despite activation and expansion within tissues. In addition, we observed higher plasma viremia in FTY720-treated animals during ATI compared to controls, particularly for the group treated also with αPD-1 and N-803. Together these results imply that tissue-localized CD8+ T cells cannot delay or prevent plasma viral rebound during ATI and suggest a direct role for vascular-restricted cytotoxic T cell subsets in control of HIV/SIV.

## RESULTS

### FTY720-mediated tissue redistribution of recirculating T cells in SIV-infected RMs

To assess the ability of tissue homing and resident CD8+ T cells to directly control viral replication in tissues, we conducted an ART interruption (ATI) study in the context of FTY720 administration after barcoded SIVmac239M infection in n=14 rhesus macaques (Fig.1A). Peripheral blood, LN samples and rectal biopsies (RB) were collected at predetermined time points during the study, with additional LN biopsies at dynamic time points immediately following viral rebound during ATI. All animals had similar peak viral loads (10^7^-10^8^ viral copies/ml) and achieved rapid control of viremia after ART initiation at day 14 p.i (Fig. 1B). As previously shown in ART-suppressed SIV-infected RMs (33, 34), FTY720 administration led to rapid and near complete redistribution of T cells from blood into tissues, with circulating CD4+ and CD8+ T cell numbers falling rapidly and remaining low throughout the FTY720 treatment period (CD4+T cells: pre-FTY720 834±304 vs. post-FTY720 13±11; CD8+ T cells pre-FTY720 596±284 vs. post-FTY720 77.2±101; Fig. 1C, representative gating strategy shown in Supplementary Fig. 1). Using MHC class I tetramers, we further quantified the absolute numbers of circulating Mamu-A*01 restricted SIV-Gag CM9 and Tat TL8 specific CD8+ T cells, which are the most common immunodominant CD8+ T cell viral epitopes of SIV (35, 36). During ART, circulating SIV-specific cell frequencies remained low in all animals, and increased in frequency in control animals after viral reactivation during ATI. No increase in circulating SIV-specific CD8+ T cells was observed in FTY720-treated RM during ART or after ATI (Fig. 1C), suggesting SIV-specific CD8 T cells that expand in response to viral rebound were maintained in lymphoid tissues in the treated RMs.

**Fig. 1.**
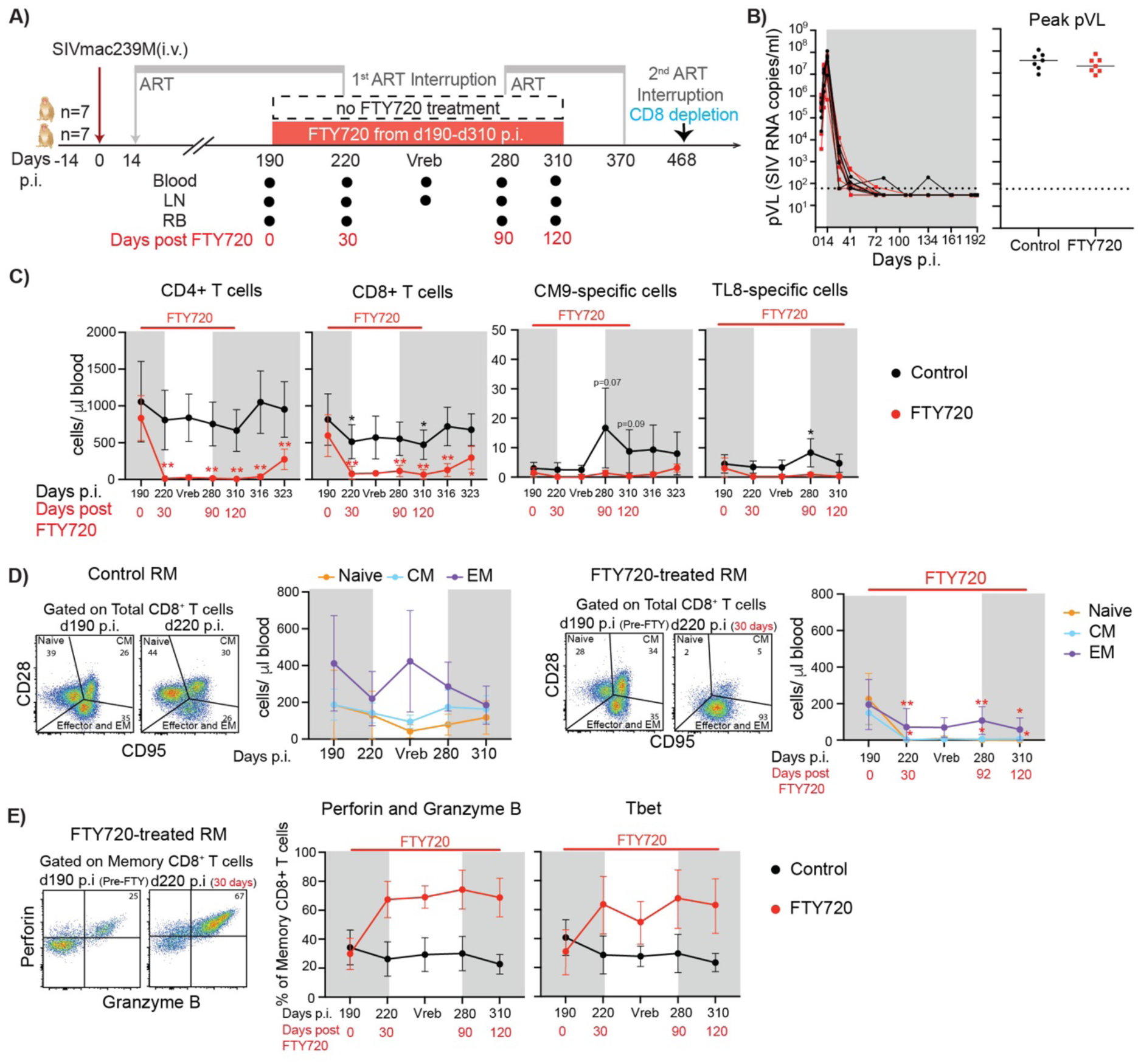
FTY720 reduces levels of circulating T cells in SIV-infected RMs during ART and at ATI. A) Schematic of the study design (i.v. = intravenously, p.i. = post infection, ART= antiretroviral treatment, pVL= plasma viral load –SIV RNA copies/ml-). B) Left: Plasma SIVmac239M RNA levels during acute infection and first six months of ART expressed as copies/ml (limit of detection = 60 copies/ml, indicated with a dashed line) are shown for each individual animal from control (black lines) or RMs treated with FTY720 from 190 to 310 days p.i. (red lines). pVL= plasma viral load (SIV RNA copies/ml). Right: peak plasma viral load during acute infection for control and FTY720-treated RMs. Lines correspond to median values. C) Absolute count of blood CD4+ T cells, CD8+ T cells, CM9- and TL8-specific CD8+ T cells shown as cells/ul of blood. D) Representative flow plots of CD28 vs. CD95 staining to evaluate memory phenotype and absolute counts of blood Naive, CM and EM CD8+ T cells of control (left) or FTY720-treated RMs (right). E) Representative flow plots of perforin versus granzyme B in FTY720-treated RM, pre and post treatment. Expression of perforin, granzyme B and Tbet on circulating memory CD8+ T cells of control or FTY720-treated RMs at different time points (n=7 for each group for every time point analyzed except at day 220 p.i. where FTY720-treated group n=5). Fresh PBMCs where used to assess CD4+, CD8+ and CM9-specific CD8+ T cell absolute counts and CD8+ T cell memory phenotype, and frozen PBMCs were used to assess TL8-specific CD8+ T cell absolute count and perforin, granzyme B and Tbet expression. Data are presented as the mean ± SD. Grey rectangles indicate time points under ART. Statistical differences in C and D were assessed by applying a 2 way-anova with Dunnett’s multiple comparison tests (every time point vs day 190 p.i.; black asterisks correspond to statistical analysis within the control arm, and red asterisks within FTY720 arm; n=7 per group per time point except viral rebound where n=3, which was excluded from the statistical analysis). *P≤0.05, **P≤0.01, ***P≤0.001, ****P≤0.0001.

While redistribution of CD4+ T cells from blood into tissues was nearly absolute in FTY720 treated animals, a population of CD8+ T cells remained in blood during FTY720 treatment. To further characterize these non-migrating CD8+ T cell populations we examined T cell memory subset distribution using CD28 and CD95 expression patterns (Fig. 1D). FTY720 strongly decreased naïve and central memory (CM) CD8+ T cell counts but had limited effect on effector/effector memory (EM) CD8+ T cells, which constituted the majority of the remaining CD8+ T cells (Fig. 1D). In FTY720-treated animals, approximately 70% of the residual circulating effector/EM CD8+ T cell populations expressed perforin and Granzyme B and 63% expressed the cytotoxicity-associated transcription factor T-bet (37) after 4 months of FTY720 administration (mean values, Fig. 1E.) We also observed an increase in the frequency of cytotoxicity-associated CX3CR1+ and a decrease in the frequency of tissue-trafficking CXCR3, CXCR5, and CCR7+ memory CD8+ T cells in FTY720 treated animals (Supplementary Fig. 2). Residual SIV-specific CD8+ T cells in blood of FTY720-treated animals also expressed an effector phenotype (data not shown).

### Expansion and activation of SIV-specific CD8+ T cells in LN of FTY720-treated RMs after ATI

To assess the impact of FTY720 on CD8+ T cells in tissues during FTY720 treatment, we measured the frequency of LN CD8+ T cells. By flow cytometry, there were no changes over time in the overall proportion of LN CD8+ T cells during FTY720 treatment compared to controls (Supplementary Fig. 3A). However, by IHC we observed an accumulation of CD8+ cells in the T cell zone of LNs of FTY720-treated RMs compared to controls after 120 days under FTY720 treatment (Fig. 2A). No change in CD8+ cell infiltration was observed in B cell follicles, nor did we detect differences between treatment groups in follicle-homing CXCR5 receptor expression by flow cytometry in memory or SIV-specific CD8+ T cells, compared to control animals (Supplementary Fig. 3B and data not shown). While CD8+ T cell memory subset distribution in the LN of control animals remained stable over time, we observed a minor increase in the proportion of EM CD8+ T cells during ATI and CM CD8+ T cells after ART reinitiation in FTY720 treated animals (Supplementary Fig 3C). Increased proliferation, as measured by Ki67, was observed during ATI in LN CD8+ T cells for FTY720-treated RMs, but no differences over time or between groups were observed for LN bulk memory or GC-TFH CD4+ T cells (Supplementary Fig. 3 D, E and F).

**Fig. 2.**
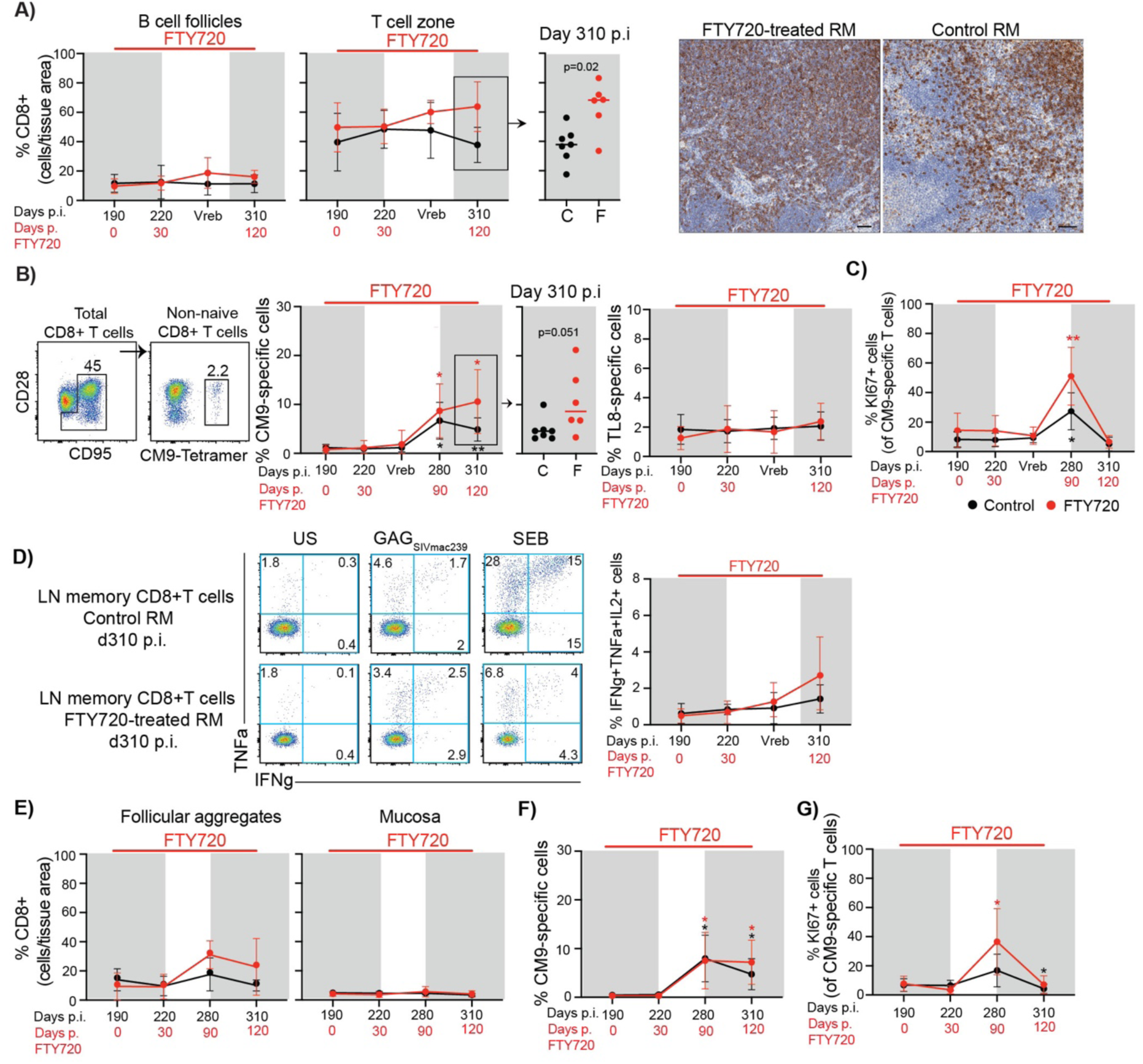
FTY720 treatment accumulated CD8+ T cells and activated CM9-specific CD8+ T cells in lymph nodes and gut mucosa during ATI. A) Lymph node CD8+ cell analysis by immunohistochemistry during FTY720 treatment (n=7 for each group at every time point, except day 190 p.i., viral rebound and day 310 p.i. for FTY720-RMs n=6). Inset showing the proportion of LN CD8+ cells per tissue area in the T cell zone at day 310 p.i., after 120 days with FTY720. Representative immunohistochemistry staining of T cell zone CD8+ cells at day 310 p.i are shown. B) Representative flow plots showing the gating strategy to identify CM9-specific CD8+ T cell in LN samples by flow cytometry. Summary plot showing the proportion of CM9-specific cells in LN of control or FTY720-treated RMs (n=7 for each group at every time point except day 280 and 310 p.i. where n=6 for FTY720-treated RMs). Inset showing the proportion of CM9-specific cells in LN of control or FTY720-treated RMs at day 310 p.i. Far right: proportion of TL8-specific cells in LN of FTY720-treated or control RM (n=7 for each group at every time point except for day 190 and 310 p.i. where n=6 for FTY720-treated RMs). C) Frequency of KI67+ cells within CM9-specific CD8+ T cells from LN of control or FTY720-treated RM (n=7 for each group at every time point, except day 280 and day 310 p.i. for FTY720-treated RMs where n=6; and day 280 p.i. for control RMs where n=6). D) Left: Representative flow plots showing TNFα versus IFNγ expression within memory CD8+ T cells after stimulation with GAG-SIVmac239 peptide pool. US = unstimulated, negative control. SEB = staphylococcus enterotoxin B, positive control. Right: Summary plots showing the proportion of CM9-specific cells either positive for IFNγ, TNFα and/or IL2 within LN memory CD8+ T cells, of control or FTY720-treated RMs over time (n=7 for each group at every time point except day 190 p.i. and day 310 p.i. where n=6 for FTY720-treated RMs). Background-subtracted values are shown. E) Rectal biopsies CD8+ T cell analysis by immunohistochemistry at different time points during FTY720 treatment (Control RMs: d190, n=5; d220: n=3; d280, n=5, d310, n=3. FTY720-treated RMs: d190, n=6; d220: n=5; d280, n=3, d310 n=2). F) Summary plot showing the proportion of CM9-specific cells in RB of control or FTY720-treated RMs over time (n=7 for each group at every time point). G) Proportion of KI67+ cells within CM9-specific CD8+ T cells of RB of control or FTY720-treated RM (n=7 for each group at every time point, except day 190 p.i. n=6 and day 220 p.i. n=5 for FTY720-treated RMs). Fresh cells from LN and RB were used for flow cytometric analysis to analyze the frequency and phenotype of CM9-specific CD8+ T cells and frozen samples were used to analyze the proportion of TL8-specific cells and cytokine production by LN SIV-specific CD8+ T cells. Grey rectangles indicate time points under antiretroviral treatment. Data are presented as the mean ± SD except when comparing between arms at a given time point where lines correspond to median values. Statistical differences were assessed by applying a 2 way-anova or mixed-effects model (if there were missing values) with Dunnett’s multiple comparison test for comparisons between time points within each arm (every time point vs day 190 p.i.; when plotted on the same graph, black asterisks correspond to statistical analysis within the control arm, and red asterisks within FTY720 arm). Mann Whitney test was used to compare between arms at day 310 p.i (A and B insets). *P≤0.05, **P≤0.01, ***P≤0.001, ****P≤0.0001.

We further analyzed the frequency and phenotype of LN SIV-specific CD8+ T cells during FTY720 treatment by MHC class I tetramer staining and cytokine production. After viral reactivation during ATI, LN Gag CM9-specific CD8+ T cells increased in frequency for both control and FTY720-treated RMs and both remained elevated after ART reinitiation, with FTY720 treated animals trending higher (Fig 2B: day 310 p.i., control vs FTY720 treated animals p=0.051). In contrast, the proportion of LN Tat TL8-specific CD8+ T cells remained steady in both animal groups (Fig. 2B). Increased proliferation of LN Gag CM9-specific CD8+ T cells was observed in both animal groups during ATI, but at significantly higher levels in the FTY720 treated group (Fig. 2C). No differences were observed between groups regarding the expression of canonical exhaustion markers (PD1, CTLA4 and TIGIT, data not shown) on SIV-specific CD8+ T cells. LN CM9-specific CD8+ T cells also produced cytokines after *ex vivo* stimulation with GAG_SIVmac239_ peptides at similar levels in both animal groups (Fig. 2D). Together, these data demonstrate that SIV-specific CD8+ T cells were present in the LN of FTY720 treated animals, and showed increased frequency and activation compared to controls while retaining functional properties.

We further addressed the effects of FTY720 on SIV-specific CD8+ T cells in mucosal sites. We observed an increase in the proportion of total and EM CD8+ T cells in rectal tissue during ATI in both treatment groups (Supplementary Fig. 3G and H). We also found a trend towards higher proportion of CD8+ T cells during ATI and after ART reinitiation in rectal tissue follicular aggregates but not mucosa of FTY720-treated RMs (Fig. 2E); however, we could not conduct statistical analysis due to several missing samples. Similar to LN, the proportion of CM9-specific CD8+ T cells increased in rectal tissue during ATI for both treatment groups (Fig. 2F), with heightened proliferation selectively observed in the FTY720 treated group during the ATI (Fig. 2G). Flow cytometric analysis of bulk memory CD8+ T cells in RB showed a similar phenotype between control and FTY720-treated RMs during ATI (data not shown).

### FTY720 induces retention of CD8+ T cells in viremic tissues but does not enable the acquisition of cytotoxic properties

Next, we examined whether tissue retention of T cells by FTY720 during ATI increased the cytotoxic properties of CD8+ T cells in lymphoid and mucosal sites. Perforin expression by LN total memory or CM9- and TL8-specific CD8+ T cells remained low and similar between control and FTY720-treated RMs, even after viral rebound during ATI (Fig. 3A), with no differential increase of perforin or granzyme B+ cells in the B cell follicles and T cell zone (Fig. 3B). We further measured total cytotoxic potential during ATI using redirected caspase-3 based killing assays and observed comparably low cytotoxic activity for LN CD8+ T cells within control and FTY720 treated groups (Fig. 3C).

**Fig. 3.**
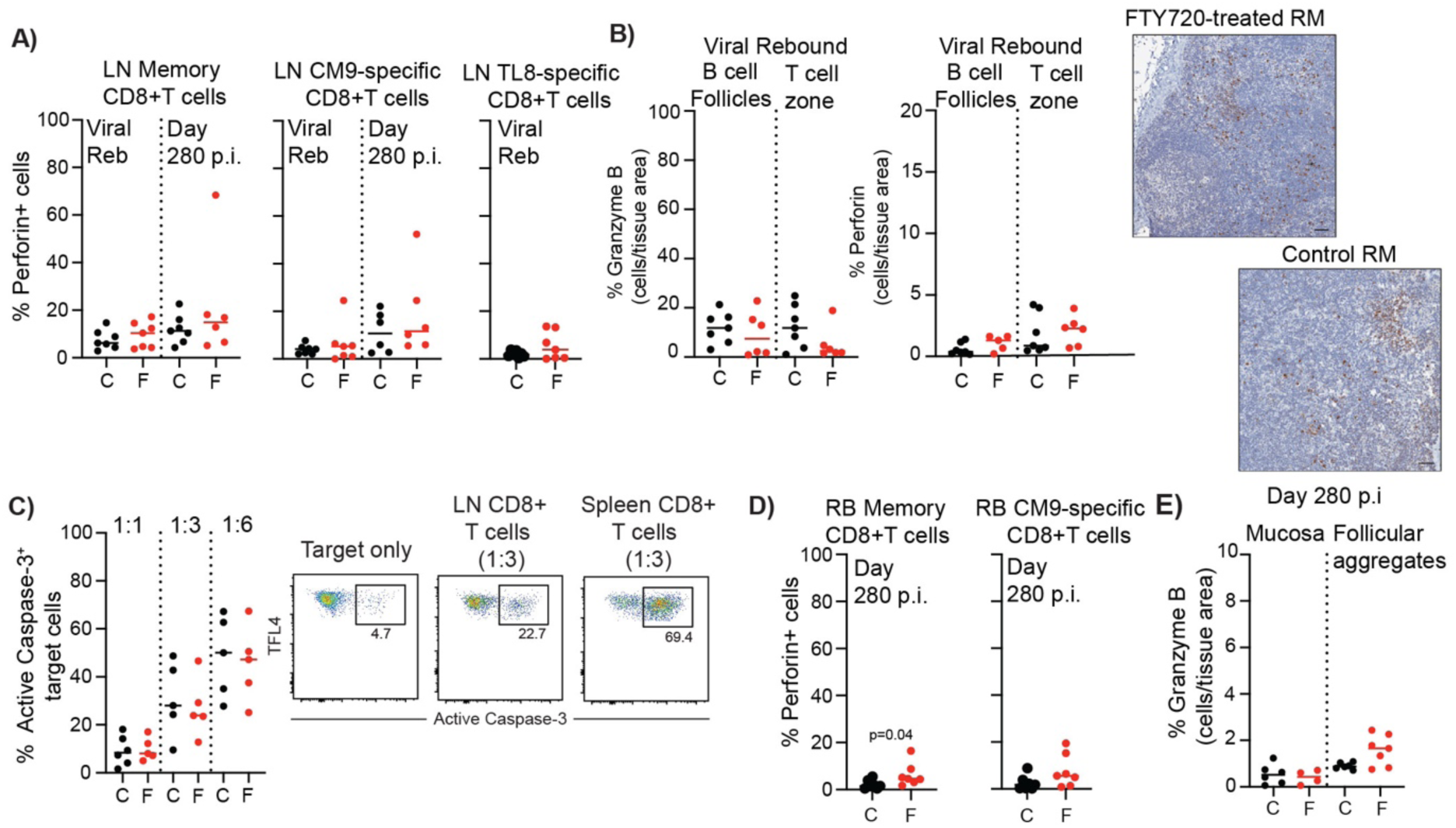
FTY720-enforced tissue retention does not enable acquisition of cytotoxic properties on CD8+ T cells. A) Perforin expression analysis of bulk memory, CM9-or TL8-specific CD8+ T cells of LN during ATI#1. Fresh cells from LN were used for flow cytometric analysis of perforin expression except for TL8-specific CD8+ T cells analysis where frozen cells were used. LN memory and CM9-specific CD8+ T cell analysis: n=7 for each group at every time point except for day 280 where n=6 for FTY720-treated RMs. LN TL8-specific cells analysis: n=7 for each group. B) Immunohistochemistry analysis of granzyme B and perforin expression on LN at viral rebound (Viral Reb) during ATI#1. Far right: Representative immunohistochemistry staining of granzyme B in LN during ATI. C) Data quantification of redirected killing assays at different target:effector ratios for LN CD8+ T cells from control and FTY720-treated RM<u>s</u> during ATI#1 (at viral rebound). Data were pregated on single live TFL4+ cells. N=5 for each group. Frozen LNMCs where used to assess T cell functionality. Representative dot plots of active Caspase-3 staining under different conditions are shown. The numbers represent the proportion of TFL4+ target active caspase-3+ apoptotic cells within single live TFL4+ cells. D) Perforin expression analysis of bulk memory and CM9-specific CD8+ T cells of rectal mucosa during ATI#1. Fresh cells from RB were used for flow cytometric analysis. N=7 for each group. E) Immunohistochemistry analysis of granzyme B and perforin expression on RB at day 280 post infection during ATI#1. Grey rectangles indicate time points under ART. Lines in the plots correspond to median values. Statistical differences were assessed with Mann-Whitney test. *P≤0.05, **P≤0.01, ***P≤0.001, ****P≤0.0001.

Rectal biopsy analysis showed similar results, with a slightly higher proportion of perforin+ memory, but not Gag CM9-specific, CD8+ T cells in FTY720-treated compared to control RMs during ATI (Fig. 3D and E).

### Inhibition of lymphocyte migration slightly limits control of viremia during ATI

FTY720 during ART was safe and well tolerated, and no viral reactivation was observed (Fig. 4A). However, following ATI all animals rebounded at similar kinetics irrespective of FTY720 treatment (Fig 4A and B; Control: 12,9±3.5 days; FTY720: 14,9±5.3 days). Furthermore, while the peak pVL and area under the curve (AUC) during the entire initial ATI were similar between control and FTY720-treated RMs (Supplementary Fig. 4A and B), control animals more effectively controlled peak viremia during the first ATI relative to acute phase peak pVL and had a lower viral set point (Fig 4C and D). To assess the potential longer-term impact of FTY720 during treatment interruption on immune control and viral reservoir stability, we reinitiated ART in all 14 animals at 2 months post ATI #1, reducing pVLs to undetectable levels within one week of ART re-initiation (Fig. 4A). Daily dosing of FTY720 was continued for one month following the re-initiation of ART in the 7 animals assigned to the treatment arm and ART was interrupted for a second time 3 months after being re-initiated. Following this second ART interruption, two animals (one from each group) exhibited post-treatment control, but otherwise no delay in viral rebound was observed in animals previously treated with FTY720 compared to control (Fig. 4E). Two animals from each group were sacrificed as soon as they rebounded. The rest of the RMs underwent experimental depletion of CD8+ T cells with an anti-CD8 antibody and, as expected, all animals demonstrated substantial viral load increases between 4 to 8 days after CD8+ T cell depletion, showing the relevance of CD8+ T cells in the control of SIV infection (Fig. 4E). Previous FTY720 treatment during the first ATI did not impact viral rebound levels or kinetics during the second ATI (Fig. 4F, Supplementary Fig. 4C, D, E), and the reduction in the peak plasma viral load compared to acute infection was similar between arms (Fig. 4G).

**Fig. 4.**
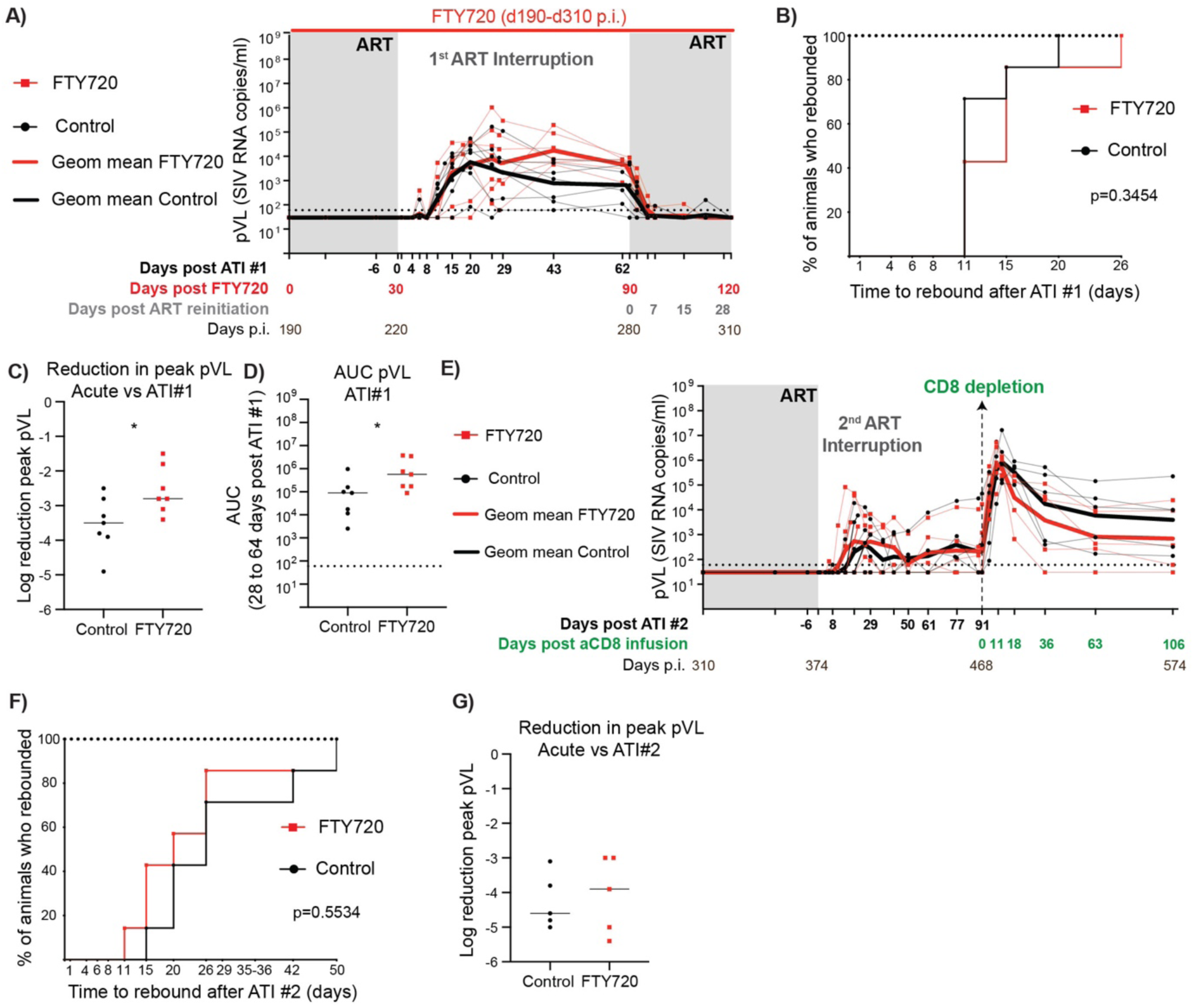
FTY720 administration did not show any therapeutic benefit on SIV infection. A) Plasma SIVmac239M RNA levels during ATI #1. Thick lines correspond to geometric mean values for each arm. B) Survival curve showing time to viral rebound during ATI #1 for control and FTY720-treated animals. C) Log reduction of peak plasma viral load during ATI#1 compared to acute infection. D) Area under the curve (AUC) from day 28 to day 64 post ATI. Lines correspond to median values. E) Plasma SIVmac239M RNA levels during ATI #2 and after CD8+ T cell depletion with anti-CD8a antibody (two animals per group were sacrificed as soon as they rebounded and anti-CD8a antibody was administered to 5 animals per group). Thick lines correspond to geometric mean values for each arm. E) Survival curve showing time to viral rebound during ATI #2 for control and FTY720-treated animals. F) Log reduction of peak plasma viral load during ATI#2 compared to acute infection. Lines correspond to median values. Statistical differences were calculated with Log-rank (Mantel-Cox) test (survival curves) and Mann-Whitney (peak pVL comparisons). *P≤0.05, **P≤0.01, ***P≤0.001, ****P≤0.0001.

### CD8+ T cell immunomodulation during FTY720 treatment is insufficient to enable viral control during ATI

Previous studies have shown that reversal of immune exhaustion or enhancement of immune function via PD1 blockade (38, 39) or IL-15R SA N-803 (40), respectively, can improve T cell responses and viral control in HIV/SIV infection. However, these immunomodulatory agents have not been used in the context of FTY720; thus, whether their effects are mediated in tissues or the periphery are unclear. To address this, we infected seven additional RMs with 10,000 IU of barcoded SIVmac239M i.v., treated with ART at d14 p.i. for 6 months, and then administered FTY720 plus N-803 and a rhesusized anti-PD1 (FNP arm) prior to and during ATI (Fig. 5A). The study design mirrored the control and FTY720-treated RM arms to be able to compare between the three treatment groups. Peak plasma viral load during acute infection was similar amongst all animals within and between the different study arms, and all animals maintained undetectable plasma viral loads during ART (Fig. 5B-C). One month prior to ATI, all animals began receiving daily doses of FTY720. Three weeks pre-ATI, they received one dose of N-803 and αPD1, with additional doses of each administered throughout the study as depicted in Figure 5A. FTY720 + N-803 + αPD1 treatment was demonstrated to be safe and well tolerated with the exception of a few instances of dermatitis and N-803 injection-site reactions. As expected, all seven RMs showed profound lymphopenia during FTY720 treatment (Fig 5D), and the majority of the remaining CD8+ T cells in blood were EM with a cytotoxic profile, similar to what was observed for FTY720-only-treated RMs (Fig 5E and Supplementary Fig. 5A and B). Additionally, PD1 receptor occupancy remained high after αPD1 mAb administration in most animals in both CD8+ T cells, preventing the detection of the receptor by flow cytometry (Fig 5F and G). Next, we evaluated the short-term effect of N-803 and αPD1 treatment on effector cells by measuring the level of proliferation of peripheral blood CD8+ T cells at several time points within 7 days post administration. We observed a transient increase in the proportion of Ki67+ SIV-specific and bulk memory CD8+ T cells post N-803 administration (Fig. 5H). Interestingly, we also observed a trend towards a transient short-term increase in the proportion of proliferating blood CD4+ T cells after N-803 and aPD1 administration (Supplementary Fig 6A). Additionally, a trend towards a minor increase in the proportion of proliferating memory CD4+T cells was observed in the LNs of FNP-treated RMs two weeks after the first doses of N-803 plus aPD1 administration during ART (Supplementary Fig 6B). This trend was not observed in LN memory CD8+ T cells at the same time point, although some animals showed an increase in the proportion of proliferating LN CD8+T cells (Supplementary Fig 6C). GC-TFH CD4+ T cells also expanded during FTY720+αPD1 and N-803 administration, prior to ATI (Supplementary Fig 6D). Furthermore, at viral rebound during ATI, the proportion of proliferating LN memory and GC-TFH CD4+ T cells was higher in FNP-treated RM compared to control and FTY-720-only treated RMs (Supplementary Fig 6E and F).

**Fig. 5.**
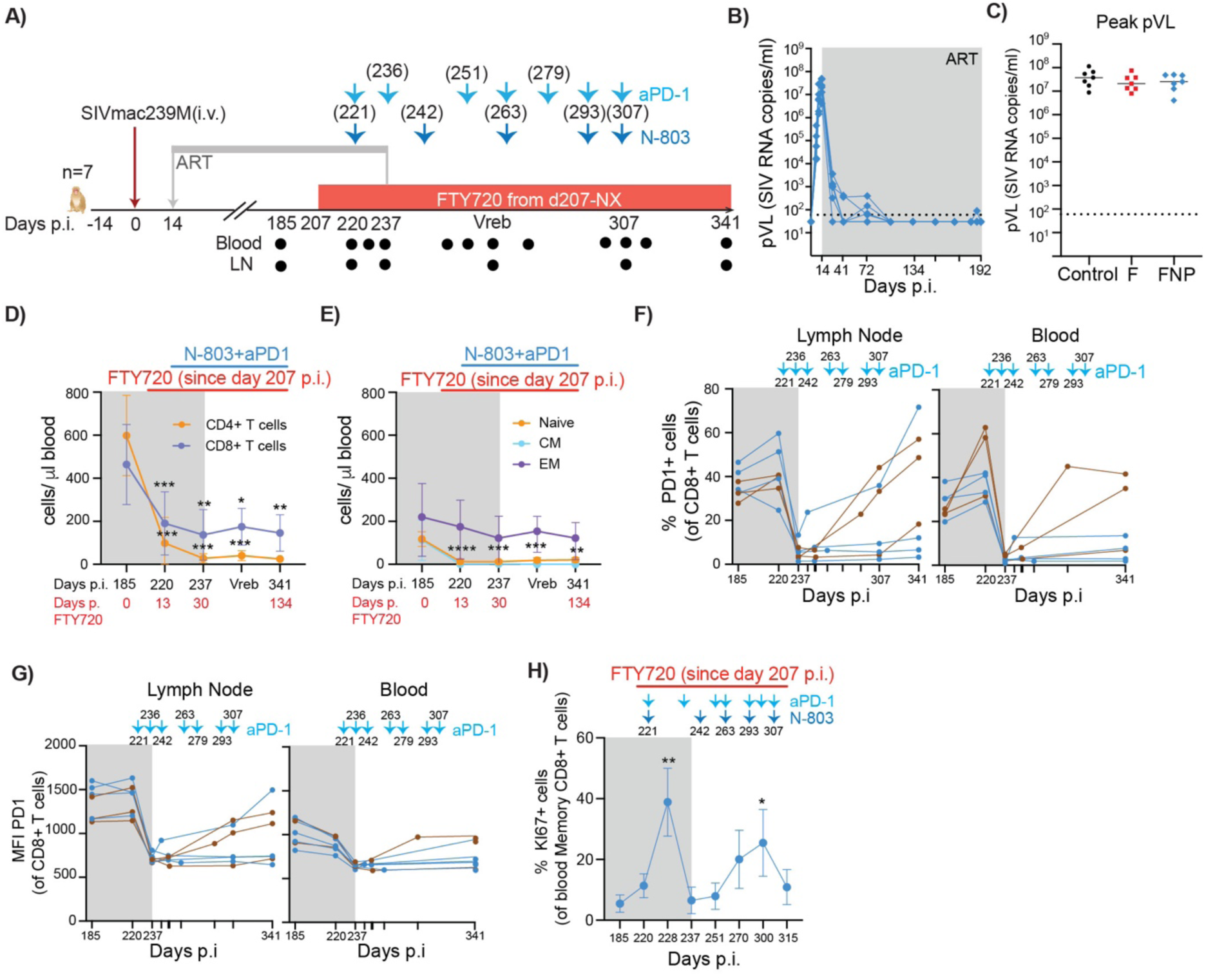
Redistribution of circulating T cells into tissues and proliferation of blood effector cells during combined FTY720, N-803 and anti-PD1 administration. A) Schematic of the study design (i.v. = intravenously, p.i. = post infection, ART= antiretroviral treatment). Numbers on top of the arrows (days p.i.) correspond to time points where aPD1 and/or N-803 were administered. Numbers (days p.i.) below the timeline correspond to time points where blood and/or LN samples were taken for mononuclear cell analysis. B) Plasma SIVmac239M RNA levels (pVL) during acute infection and first six months of ART expressed as copies/ml (limit of detection = 60 copies/ml, indicated with a dashed line) are shown for each individual animal. C) Summary plot showing the peak plasma viral load during acute infection for control, FTY720 and FTY720-aPD1-N-803-treated RMs. Lines correspond to median values. D) Absolute count of blood CD4+ T cells and CD8+ T cells and E) absolute count of naïve, central memory (CM) and effector memory (EM) CD8+ T cells in FTY720-N-803-aPD1-treated RMs at selected time points pre and post FTY720-administration. F) and G) Summary plots showing PD1 receptor occupancy (proportion of positive cells and mean fluorescence intensity, MFI) in LN mononuclear cells and PBMCs CD8+ T cells over time. Arrows indicate aPD1 administration and numbers on top or below the arrows are days p.i. H) Summary plot showing the changes in KI67 (surrogate for proliferation) in peripheral memory CD8+ T cells over time. Arrows indicate aPD1 and/or N-803 administration. Numbers below the arrows are days p.i. Mean and standard deviation values are shown. Cryopreserved mononuclear cells were used to assess T cell phenotype. Grey rectangles indicate ART. Statistical differences in C) were calculated using Kruskal-wallis test with Dunn’s post comparisons versus control group; in D), E) and H) one-way anova with Dunnet’s post comparisons every time point vs day 185 p.i (D and E) or vs day 220 (H). *P≤0.05, **P≤0.01, ***P≤0.001, ****P≤0.0001.

Following ATI, all animals rebounded, with three animals (Fig. 6A, brown lines) achieving intermittent post-ATI control of viremia. Despite these animals demonstrating control, the FNP combination did not otherwise yield a beneficial effect, as there was no delay in viral rebound and we observed a trend towards higher peak viral loads compared to the control group (Fig. 6B-D). Furthermore, not only the reduction in peak pVL during ATI compared to acute infection was larger for control compared to FNP-treated RMs but also the viral set point during ATI was higher for FNP-treated RMs, similar to that observed when comparing control vs only FTY720-treated animals (Fig. 6E and F).

**Fig. 6.**
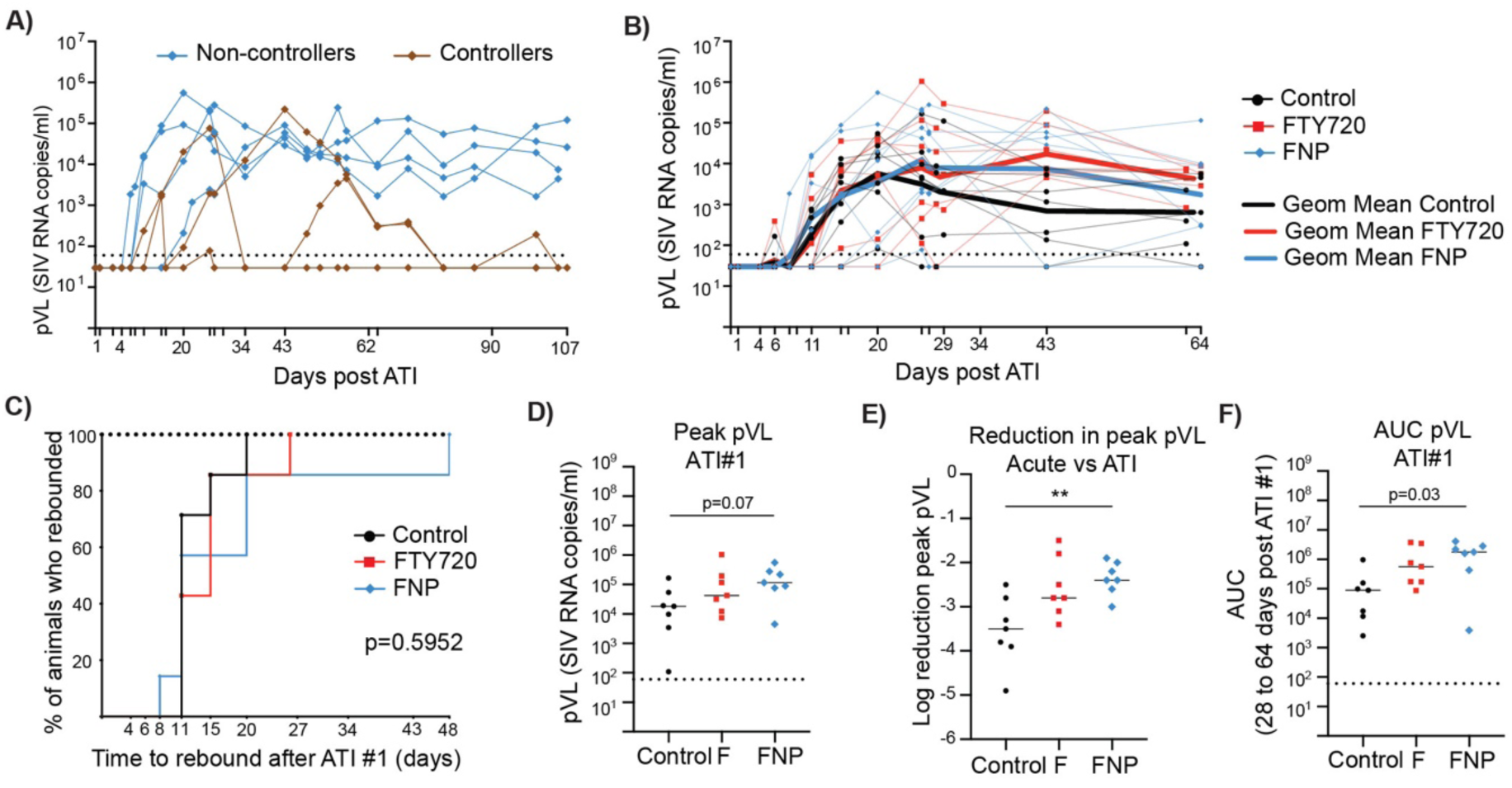
FTY720, N-803 and anti-PD1 administration impacts plasma viral load during ATI. A) Plasma SIVmac239M RNA (pVL) levels during ATI are expressed as copies/ml (limit of detection = 60 copies/ml, indicated with a dashed line). 3/7 animals that controlled the infection during ATI are shown in brown. B) Plasma SIVmac239M RNA levels during ATI#1 for control and FTY720-treated RMs. For comparison purposes, pVL from equivalent timepoints of FTY720-N-803-aPD1-treated RMs were included in the summary plot. Thick lines correspond to the geometric mean plasma viral load levels for each group. C) Survival curves showing time to viral rebound during ATI #1 for control, FTY720 and FTY720-N-803-aPD1-treated animals. D) Summary plot showing the peak plasma viral load during ATI#1 for control, FTY720 and FTY720-aPD1-N-803-treated RMs. E) Log reduction of peak plasma viral load during ATI#1 compared to acute infection for each treatment group. F) Area under the curve (AUC) from day 28 to day 64 post ATI. Grey rectangles indicate ART. Statistical differences in C) were calculated with Log-rank (Mantel-Cox) test; and in D), E) and F), using Kruskal-wallis test with Dunn’s post comparisons versus control group. *P≤0.05, **P≤0.01, ***P≤0.001, ****P≤0.0001.

Finally, we evaluated if αPD1 and N-803 treatment combined with FTY720-enforced retention increased the cytotoxic ability of LN CD8+ T cells using samples collected at various stages during the treatment period (Fig. 5A). While we observed an increase in the frequency of LN SIV-specific CD8+ T cells during ATI (Fig 7A), the proportion of LN cytotoxic SIV-specific CD8+ T cells remained low and constant over time by flow cytometry and imaging analysis (Fig. 7B and C). We also did not observe an increase in cytotoxic ability of total LN CD8+ T cells in redirected killing assays (Fig. 7D).

**Fig. 7.**
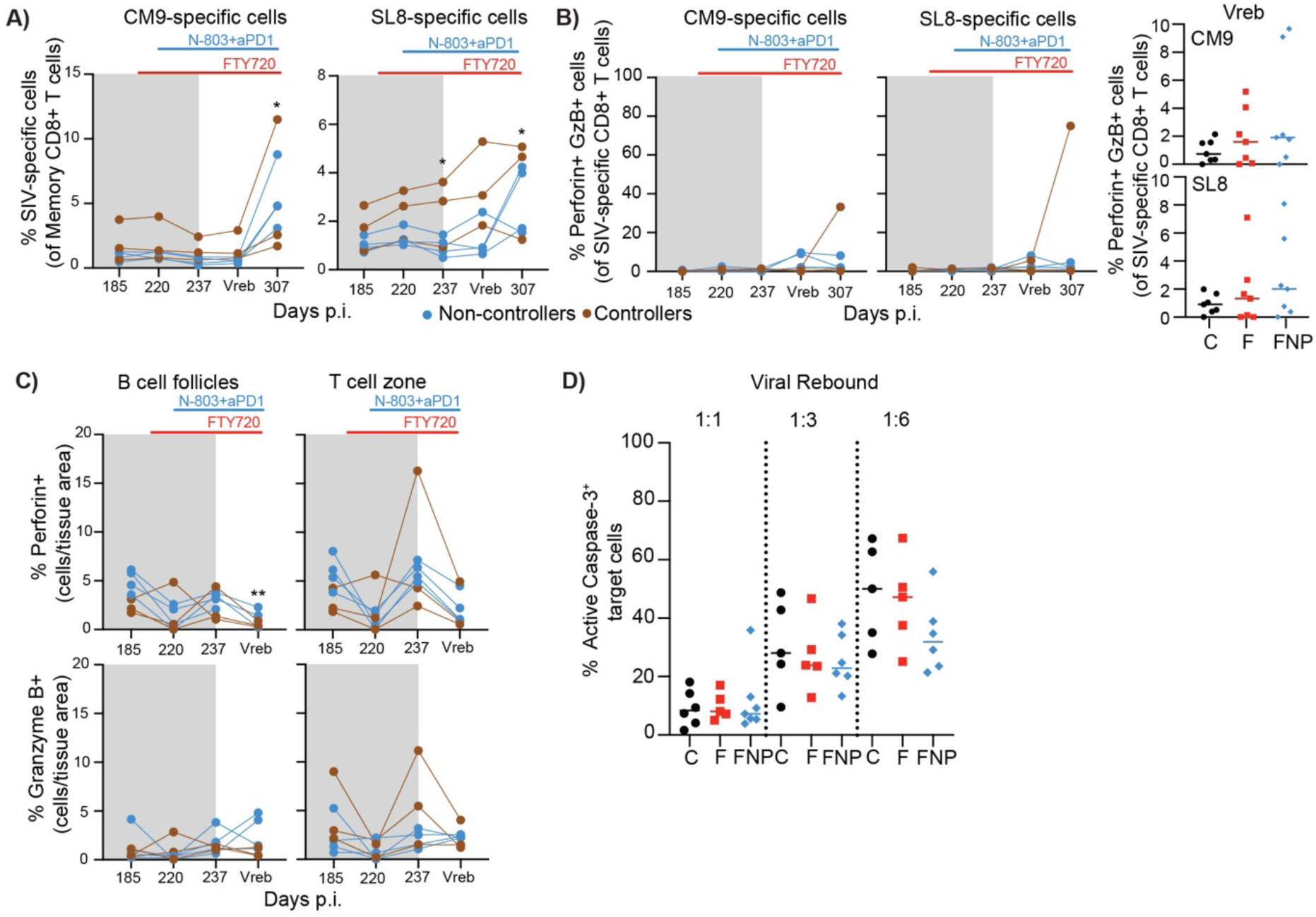
N-803 and aPD1 administration during FTY720-enforced tissue retention does not enable acquisition of cytotoxic properties on CD8+ T cells. A) Summary plot showing the proportion of LN CM9- and SL8-specific CD8+ T cells over time (n=7). 3/7 animals that showed intermittent control of the infection during ATI are depicted in brown. B) Summary plots showing the proportion of perforin and granzyme B double positive cells within LN CM9- or SL8-specific cells over time, analyzed by flow cytometry (n=7). Inset: Proportion of cytotoxic cells within LN SIV-specific CD8+ T cells at viral rebound of each arm (control, FTY720- or FTY720-N-803-aPD1-treated RMs; n=7 for each group). Lines correspond to median values. C) Immunohistochemistry analysis of granzyme B and perforin expression on LN over time in FTY720-N-803-aPD1-treated RMs (n=7, except for perforin analysis on B cell follicles at viral rebound where n=6). D) Data quantification of redirected killing assays at different target:effector ratios for LN CD8+ T cells from control, FTY- or FTY720-N-803-aPD1 treated RMs at viral rebound. Data were pre-gated on single live TFL4+ cells. N=7 for each group. Lines correspond to median values. Frozen LNMCs where used to assess T cell phenotype and functionality. Grey rectangles indicate time points under ART. Statistical differences were assessed by applying a 2 way-anova or mixed-effects model (if there were missing values) with Dunnett’s for comparisons between time points within each arm (every time point vs day 185 p.i.). Differences between groups were calculated using Kruskal-wallis test with Dunn’s post comparisons versus control group. *P≤0.05, **P≤0.01, ***P≤0.001, ****P≤0.0001.

## DISCUSSION

While the role of HIV/SIV-specific CD8+ T cells in control of HIV/SIV is well established (8, 19–21), CD8+ T cells remain unable to fully eliminate the HIV/SIV reservoir, resulting in chronic viremia and disease progression in the absence of antiretroviral therapy. This shortcoming is due in part to impaired cytotoxic CD8+ T cell activity in lymphoid tissues in which these cells are primarily cytokine producers and fail to mature into cytotoxic effectors (26, 27, 41, 42), and the differential LN migration capabilities of early differentiated (naïve and central memory) vs. late differentiated (cytotoxic effector) CD8+ T cells in the vasculature. While each of these mechanisms normally help to protect the host from immune-mediated destruction of antigen presenting cells, in the case of HIV/SIV infection, migration of infected CD4+ T cells into lymphoid tissues potentially allows evasion of peripheral cytotoxic CD8+ T cell surveillance. Even for lymphoid CD8 T cell subsets correlating with better viral control, such as those expressing both TCF-1 and CD39, their antiviral role seems largely independent from canonical cytotoxic functions (31).

Here, we tested this concept using FTY720 to inhibit T cell tissue egress in SIV-infected RM, resulting in physical separation of the SIV-infected CD4+ T cells from cytotoxic SIV-specific CD8+ T cells in blood. In this scenario, the only CD8+ T cells that could encounter, and potentially eliminate, SIV-infected CD4+ T cells after ATI would be those trapped within the same tissues. We found that FTY720 was highly effective at preventing T cell migration in SIV-infected RM, with nearly all T cells remaining in tissues except for late-differentiated cytotoxic CD4+ and CD8+ T cells. Importantly, LN and mucosal CD8+ T cells became activated after ATI, indicating target cell recognition, but did not acquire cytotoxic ability. We observed no viral blips or T cell activation during FTY720 prior to ATI, indicating that general immune quiescence persists during this redistribution event. Nevertheless, following ATI, FTY720-treated animals demonstrated viral rebound with similar kinetics to controls and trended towards higher peripheral blood viremia, especially in animals treated with αPD-1+N-803 in combination with FTY720. Previous publications have shown a modest impact on viral control after FTY720 administration, at least in a subset of SIV-infected RM (33, 34), however, FTY720 was administered during ART in these studies, not during ATI.

Based upon the near absence of CD4+ T cells in the blood of the FTY720-treated RMs, plasma viremia in these animals must originate from infected cells in tissues, and directly implicates either poor immunosurveillance or no immune-mediated clearance in these sites. Normally, infected CD4+ T cells migrate between the vasculature and tissues, subjecting a portion of the infected cells to vascular immune surveillance. However, in the FTY720 treated animals, nearly all infected cells became trapped in tissues, and thus were not subject to vascular surveillance. Alternatively, sequestration into the tissues may have provided activation or proliferative cues to the CD4+ T cells, leading to increased viral replication. However, we did not observe increased activation or cell cycle entry (Ki67) of bulk LN CD4+ T cells or GC-TFH CD4+ T cells after FTY720 administration prior to or after ATI. These results indicate that LT-CD8+ T cells alone are insufficient to delay or prevent plasma viral rebound during ATI, and further suggest that the absence of cytotoxic CD8+ T cell activity in tissues plus the physical separation of SIV-infected CD4+ T cells from the cytotoxic blood CD8+ T cells were the primary driver of the increased viral load in FTY720-only treated animals. We did indeed observe an increase in the proportion of proliferating CD4+ T cells and GC-TFH in LN of FNP-treated animals. Immunomodulation with αPD-1 or IL15R SA have individually been shown to induce CD4+ T cell activation (38, 39, 43, 44); thus, it is possible that the heightened viral load in the RM treated with FTY720 and αPD-1+ N-803 resulted in part from increased viral replication or viral target cell availability.

These results have significant impact towards the development of viable strategies to engage CD8+ T cell mediated immune clearance of tissue viral reservoirs. First, our findings suggest that vascular-restricted cytotoxic CD8+ T cells play a direct role in control of viremia but require interaction with infected CD4+ T cells for their elimination within the vascular space. Infected CD4s in tissues must migrate into the blood for clearance through this mechanism, and strategies designed to mobilize infected cells into circulation may enhance reservoir clearance. Second, simply expanding the number of CD8+ T cells in LT does not enable control of viral replication after ATI, because tissue trafficking and resident T cells cannot efficiently eliminate infected cells due to poor cytotoxic ability. Furthermore, memory CD8+ T cells within lymphoid and mucosal tissues are unable to acquire cytotoxic functions after exposure to renewed viral replication within the tissue. The reasons for this are currently unclear but may be due to both cell intrinsic and cell extrinsic regulatory effects. For example, TGFβ may negatively modulate cytotoxic properties in LT CD8+ T cells, as it was demonstrated that the in vitro neutralization of TGFβ partially restored perforin expression in gut CD8+ T cells(41). Finally, despite evidence of PD1 blockade and IL15R SA bioactivity within peripheral blood and lymphoid tissues, we did not observe increased CD8+ T cell cytotoxicity in tissues or evidence of viral control. These findings contrast with previous publications (38–40, 43), likely because FTY720 administration during the immunotherapy phase prevented the interaction between SIV-infected CD4+ T cells and cytotoxic SIV-specific CD8+ T cells in blood. We also did not find evidence of T cell exhaustion in tissues, as SIV-specific CD8+ T cells in LN of FTY720 treated RM became activated and expanded *in vivo* and produced cytokines upon stimulation in vitro. This further reinforces a previously unappreciated role for T cell trafficking and anatomical differences in T cell function in the control of HIV/SIV disease progression.

HIV cure-based strategies designed to engage CD8+ T cell-mediated elimination of HIV reservoirs in tissues have long been in development, including therapeutic vaccine, viral reactivation, exhaustion reversal, and chimeric antigen receptor (CAR) based modalities (45, 46). While these strategies have shown promise in vitro or in small animal models, immunological clearance of tissue reservoirs in HIV/SIV infection has thus far remained elusive. Here we have identified a critical issue that undermines the potential of these strategies, specifically that SIV-specific CD8+ T cells in LN and gut tissues of SIV-infected RM do not become cytotoxic after interaction with virally infected cells, even when provided optimal immunotherapy and prolonged exposure. As such, our results suggest that engagement of CD8+ T cell cytotoxic activity for eradication of tissue HIV reservoirs may require mobilization and reactivation of tissue-localized infected CD4+ T cells into the peripheral blood, or modification of the trafficking ability of cytotoxic CD8+ T cells in the peripheral blood to enable entry into tissue reservoir sites. These strategies, in combination with potent latency reversal agents, exhaustion reversal and therapeutic vaccine-mediated expansion, may enable CD8+ T cell mediated clearance of the HIV reservoir.

## MATERIALS AND METHODS

### Animal Models and Study Design

Indian-origin Rhesus Macaques (RM, *Macaca mulatta*) were housed in an animal biosafety level 2 (BSL-2) facility at the Emory National Primate Research Center in Atlanta, Georgia as previously described (47).

For the first two arms of this study, 14 female Mamu A*01 (average age of 6 years and 1 month) were infected intravenously with 10,000 copies of SIVmac239M, a genetically tagged virus with a 34-base genetic barcode inserted between the vpx and vpr accessory genes of the infectious molecular clone SIVmac239 (48). Some aspects of this study were previously reported elsewhere (18). At day 14 post-infection (p.i.) all animals were placed on ART [Tenofovir Disoproxil Fumarate (TDF; 5.1 mg/Kg per day), emtricitabine (FTC; 40 mg/Kg per day) and dolutegravir (DTG; 2.5 mg/Kg per day)] formulated in a single daily injection (1ml/Kg per day; s.c.) (Figure 1A). ART was continued for approximately 6 months (up to day 220 p.i.). One month prior to ART cessation, macaques were stratified into FTY720 only vs. control groups based on peak plasma viral load during acute infection, age, weight, and time taken to suppress viremia on ART. Thirty days prior to ATI (at day 190 p.i.), seven animals were placed on FTY720 (orally, 500 μg/kg/day) for a total of four months (until day 310 p.i.). At day 280 p.i., ART was reinstated for three additional months until a second ATI (day 373 p.i.). After viral rebound, 4 RM were necropsied, and the remaining 10 RM received MT807R1 CD8a depletion antibody (subcutaneous, 50 mg/kg) at day 468. The remaining RM were necropsied by day 574 p.i.

In the third arm of the study, a cohort of 7 Mamu A*01 male Indian-origin RMs (average age of 4 years and 11 months) were infected intravenously with 10,000 copies of SIVmac239M 3 years after the first two arms and treated daily beginning at day 14 p.i. with the previously described ART regimen. ART was continued for approximately 6 months and interrupted at day 236 p.i.. One month prior to ART cessation (day 207 p.i.), animals were placed on FTY720 (orally, 500 μg/kg/day) which was continued until necropsy at day 341/343 p.i. 15 days before ATI (day 221 p.i.), animals received their first of 5 doses of N-803 IL-15R SA (subcutaneous, 100 ug/kg) and their first of 7 doses of αPD1 (intravenous, 3mg/kg). Animals received the remaining 4 doses of N-803 at an interval of 2-4 weeks on days 242, 263, 293 and 307 p.i. and the remaining 6 doses of αPD1 at ART interruption (day 236 p.i.) and days 251, 263, 279, 293 and 307 days p.i..

### FTY720, MT807R1, N-803, and αPD-1 Administration

FTY720 (Sigma-Aldrich, CAS # 162359-56-0) was reconstituted in HPLC water to a concentration of 12.5mg/mL and administered orally at a dose of 500ug/kg with food unless animals were anesthetized for another procedure, in which case FTY720 was delivered via an orogastric feeding tube. The rhesus IgG1 recombinant Anti-CD8 alpha [MT807R1] monoclonal antibody was obtained from the Nonhuman Primate Reagent Resource (NIH Nonhuman Primate Reagent Resource Cat# PR-0817, RRID:AB_2716320), and administered at a concentration of 10.6 mg/ml and dose of 50 mg/kg subcutaneously across multiple sites (∼10-12 ml/site). N-803 was provided by NantKwest/Immunity Bio as a pharmaceutical-grade compound and administered at a concentration of 2mg/mL and a dose of 100 ug/kg subcutaneously across 1-4 locations on the upper and midback. Rhesusized αPD-1 (NIVOR4LALA) was provided by the Nonhuman Primate Reagent Resource and formulated as 10.15 mg/ml in 20 mM sodium citrate, 50 mM NacL, 3% MALTOSE, pH 6, 0.02%. αPD-1 was administered intravenously at a concentration of 10.15 mg/ml and a dose of 3 mg/kg.

### Sample collection and processing

Peripheral blood, LN biopsies, LN fine needle aspirates (FNA), plasma, and rectal biopsies (RBs) were collected longitudinally, including at necropsy. Peripheral blood mononuclear cells (PBMCs) were separated by Ficoll-Hypaque density centrifugation (Cytivia) and cryopreserved in 90% fetal calf serum (GeminiBio) plus 10% dimethyl sulfoxide (Sigma-Aldrich). Macaques were anaesthetized with Telazol (3-5mg/kg, IM) for surgical preparation of LN and RBs. LN and FNA samples were collected and processed as described elsewhere (33). Half of each LN biopsy was paraffin fixed for immunohistochemistry (IHC), while the other half was homogenized and passed through a 70-μm cell strainer to obtain mononuclear cells for flow cytometric analysis and cryopreservation for future analysis. RB were collected by inserting an anoscope into the rectum and collecting 20 RB punches by forceps. Part of the tissue was paraffin embedded for IHC analysis. To obtain gut-derived lymphocytes for flow cytometry analysis, these punches were collagenase treated (1mg/ml) for 2 hours at 37C on a shaker and then passed through a 70-μm cell strainer to remove the tissue residues.

### Plasma viral load quantification

Plasma SIVmac239M loads were quantified at the Virology Core of the Emory Center for AIDS Research using a standard quantitative PCR (qPCR) assay (with a limit of detection of 60 copies/ml) as described previously (49). The time to viral rebound was considered to be the first day with detectable viral load after ATI. Additionally, time to viral rebound was assessed as the first day of two consecutive days with a detectable viral load after ATI. No differences were observed between both methods. Day post ATI was calculated by defining the first day without ART as day 1 post ATI.

### Fluorescence cytometry staining

Fresh or cryopreserved PBMC and LN mononuclear cells (LNMC), and fresh gut-derived lymphocytes were used for flow cytometry analysis. Cryopreserved PBMC and LNMC were thawed, counted, examined for viability, and rested for 2 hours for tetramer staining or overnight for lymphocyte stimulation experiments at 37 °C and 5% CO2 in complete R10 media (RPMI supplemented with 10% FBS, 2 mM L-glutamine, 100 U/ml penicillin, 100 mg/ml streptomycin) with 10U/mL DNAse I (Roche Life Sciences, Indianapolis, IN). MHC-Class I tetramer staining was done at room temperature (RT), for 10 minutes. Then, cells were stained for viability exclusion using Live/Dead Fixable Aqua (Invitrogen) for 10 minutes, followed by a 20-minute incubation with a panel of directly conjugated monoclonal antibodies diluted in equal parts of fluorescence-activated cell sorting (FACS) buffer (PBS containing 0.1% sodium azide and 1% bovine serum albumin) and Brilliant stain buffer (BD Biosciences, San Jose, CA), also at RT. Cells were washed in FACS buffer (0.1% sodium azide and 1% bovine serum albumin in 1X PBS) and then fixed/permeabilized using the FoxP3 Transcription Factor Buffer Kit (eBioscience, San Diego, CA), following manufacturer’s instructions. Intracellular staining was performed by adding the antibody cocktail prepared in 1X perm-wash buffer for 1 hour at RT. Stained cells were washed and fixed in PBS containing 1% paraformaldehyde (Sigma-Aldrich, St. Luis, MO), and stored at 4°C in the dark until acquisition. Samples were acquired using a FACS Symphony A5 or LSR Fortessa cytometer within 48 hours. See table S1 for antibody information and Supplementary Figure 1 for the gating strategy.

### MHC Class I Tetramers

Tat-TL8 (TTPESANL) Mamu-A*01-APC, Tat-SL8 (STPESANL) Mamu-A*01-APC, Gag-CM9 (CTPYDINQM) Mamu-A*01-APC conjugated tetramers, and Gag-CM9 Mamu-A*01 monomers were supplied by the NIH Tetramer core facility at Emory University, Atlanta, GA. Gag-CM9 Mamu A*01-BV421 tetramers were prepared in our laboratory using Gag-CM9 monomers and streptavidin-BV421 (Biolegend, San Diego, CA). The Mamu-A*01 Tat_28-35_ tetramer was constructed using an SIV_MAC251_-derived peptide. Even though the corresponding SIV_MAC239_ sequence was STPESANL, this tetramer detected strong responses in SIV_MAC239_-infected macaques, as it has been shown before where Tat_28-35_ (STPESANL) and Tat_28-35_ (TTPESANL) tetramer staining yielded identical results (35).

### Lymphocyte stimulation

For stimulation assays, mononuclear cells were plated in a V-bottom 96-wells plate at 1.5M/200ul of R10 media. SIVmac239 Gag Peptide Pool was obtained from the NIH AIDS Reagent Program, Division of AIDS, NIAID, NIH, and the working concentration was 2 μg/ml. Co-stimulation was added with peptides: 1 μg/mL anti-CD49d (Clone 9F10, Biolegend, San Diego, CA) and CD28-ECD (Clone CD28.2, Beckman Coulter, Brea, CA). A DMSO-treated condition served as a negative control. Positive control samples were stimulated using Staphylococcal Enterotoxin B (SEB, List Biological Laboratories, Campbell, CA) at 1 μg/mL. CD107a BV650 (clone H4A3, Biolegend, San Diego, CA) was added at the start of stimulation. Brefeldin A (1μg /mL) (Sigma Aldrich, Saint Louis, MO) and monensin (0.66μL/mL) (BD Biosciences, San Jose, CA) were added one hour after initiation of stimulation. Cells were incubated under stimulation conditions for a total of 9 hours. After washing with PBS, cells were stained as described above. Samples were acquired using a BD FACS Symphony A5 flow cytometer (BD Biosciences). All flow cytometry data were analyzed using FlowJo V10.7.1 software (Tree Star, Ashland, OR) and GraphPad Prism (Version 10.2.2 GraphPad Software, La Jolla California USA). Data shown in figures have been background-subtracted.

### Immunohistochemistry

Immunohistochemical staining and quantification were performed as previously described (50). In brief, immunohistochemistry on LN biopsies was performed on 5-μm tissue sections. Heat-induced epitope retrieval was performed by heating sections in 0.01% citraconic anhydride containing 0.05% Tween-20 then incubated with antibody to Granzyme B (GrzB, HPA003418, Sigma, 1:200) or CD8 (NBP2-34039, NOVUS, 1:1000) diluted in TBS Tween overnight 4°C. Slides were washed in 1× TBS with 0.05% Tween-20, endogenous peroxidases blocked using 1.5% (v/v) H2O2 in TBS, pH 7.4, for 5 min, incubated with rabbit or mouse Polink-1 horseradish peroxidase (HRP) and developed with ImmpactTM DAB (3,3′-diaminobenzidine; Vector Laboratories) according to manufacturer’s recommendations. All slides were washed in H_2_O, counterstained with haematoxylin, mounted in Permount (Fisher Scientific), and scanned at high magnification (x200) using the ScanScope CS System (Aperio Technologies), yielding high-resolution data from the entire tissue section. Representative regions of interest (0.4 mm^2^) were identified, and high-resolution images extracted from these whole-tissue scans. The percentage area positive for GrzB positive cells was quantified using Cell profiler version 3.1.5 (51).

### Re-directed killing assay

Cytolytic activity of LN CD8+ T cells was evaluated using an adapted re-directed killing assay for non-human primates (NHP). Cryopreserved LN lymphocytes and splenocytes were thawed, counted, examined for viability, and rested overnight at 37 °C and 5% CO2 in RPMI supplemented with 10% FBS, 2 mM L-glutamine, 100 U/ml penicillin, 100 mg/ml streptomycin and 10U/ml DNAse I. CD8+ T cells were negatively selected from LN or spleen using a NHP CD8+ T Cell Enrichment Kit (StemCell Technologies). P815 mastocytoma target cells were labeled with LIVE/DEAD Fixable Violet (Thermo Fisher Scientific) and TFL4 (OncoImmun), washed twice in PBS, and incubated for 30 min at room temperature with α-CD3 (1 mg/mL; clone SP34-2; BD Biosciences). Isolated CD8+ T cells were rested in complete medium for at least 45 min at room temperature and then incubated with α-CD3-coated P815 cells at different target:effector ratios in a 96-well V-bottom plate for 4 hr at 37 °C and 5% CO_2_. Cells were washed with PBS and then fixed/permeabilized using the BD Cytofix/Cytoperm Fixation/Permeabilization Kit (BD), following manufacturer’s instructions.

Cells were then stained with α-active caspase 3 and α-CD8 prepared in 1X permwash buffer for 1 hour. Stained cells were washed and fixed in PBS containing 1% paraformaldehyde and stored at 4°C in the dark until acquisition. Cells were acquired using a BD FACS Symphony A5 flow cytometer (BD Biosciences). All flow cytometry data were analyzed using FlowJo V10.7.1 software and GraphPad Prism.

### Statistical analysis

Data analyses were performed using GraphPad Prism. Statistical significance of viral data between study groups was performed using Mann-Whitney or Kruskal-Wallis with Dunn’s multiple comparisons test vs control group when comparing three groups. Differences in time to viral rebound were assessed using the Log-rank (Mantel-Cox) test. Statistical significance of absolute cell count and immunophenotyping were assessed by applying a one or two way-anova or mixed-effects model (if there were missing values) with Dunnet’s multiple comparison test for comparisons between time points within each arm (every time point vs day 190 p.i., unless stated otherwise). For immunophenotype pairwise comparisons at a given time point, Mann-Whitney test was used to compare values between two arms, or Kruskal-Wallis with Dunn’s multiple comparisons test vs control group was used when comparing three groups. A *P* value less than 0.05 was considered statistically significant. *P≤0.05, **P≤0.01, ***P≤0.001, ****P≤0.0001.

### Study approval

All procedures were approved by Emory University Institutional Animal Care and Use Committee (IACUC) under permit PROTO201700655.

## List of Supplementary Materials

Supplementary figure S1-S6 and table S1.

## Author contributions

Conceptualization: MRB, MP, MPD, KJB

Investigation: MBP, SS, EV, CD, LKC, JR, VW, MPD, SSD

Formal analysis: MBP, EV, CD Resources: BFK, JTS Visualization: MBP, MRB

Funding acquisition: MRB, MP, MPD, KJB Supervision: MRB, MP

Writing – original draft: MBP, MRB

Review and editing: MBP, MRB, EV, MP, MPD, SSD, VW, BFK, CD, SS

## Acknowledgments

Part of the data for this manuscript were generated in the Penn Cytomics and Cell Sorting Shared Resource laboratory at the University of Pennsylvania which is partially supported by the Abramson Cancer Center NCI Grant (P30 016520). The research identifier number is RRid:SCR_022376. We thank Jennifer Wood, Sherrie M. Jean, Stephanie Ehnert, and Stacey Weissman (Emory National Primate Research Center -EPC-Division of Animal Resources and Research Resources) for providing support in animal and veterinary care. Plasma viral loads were conducted by Thomas Vanderford, Shan Liang, and Shelly Wang at the Emory Center for AIDS Research (CFAR) Virology Core. We thank Gilead for providing TDF and FTC, and ViiV Healthcare for providing DTG.

## Supplemental material

**Table S1:** Antibodies used for flow cytometry.

| Antibody | Fluorochrome | Clone | Source |
| --- | --- | --- | --- |
| Granzyme B | AF700 | GB11 | BD Bioscience |
| CXCR3 | BUV395 | IC6/CXCR3 | BD Bioscience |
| CD8 | BUV563 | RPA-T8 | BD Bioscience |
| CD8 | BUV496 | RPA-T8 | BD Bioscience |
| CD4 | BUV661 | SK3 | BD Bioscience |
| CD95 | BUV737 | DX2 | BD Bioscience |
| CD95 | BV650 | DX2 | BD Bioscience |
| CD95 | PE-Cy5 | DX2 | BD Bioscience |
| CD3 | BUV805 | SP34-2 | BD Bioscience |
| CD3 | BUV395 | SP34-2 | BD Bioscience |
| HLA-DR | BV480 | G46-6 | BD Bioscience |
| HLA-DR | BV605 | G46-6 | BD Bioscience |
| HLA-DR | BUV395 | G46-6 | BD Bioscience |
| KI67 | BV785 | B56 | BD Bioscience |
| KI67 | AF700 | B56 | BD Bioscience |
| IFNg | BV750 | B27 | BD Bioscience |
| IL-2 | PE | MQ1-17H2 | BD Bioscience |
| CCR7 | PE-Cy7 | 3D12 | BD Bioscience |
| CD16 | BUV615 | 3G8 | BD Bioscience |
| CTLA-4 | BV421 | BNI3 | BD Bioscience |
| Caspase 3 | PE | C92-605 | BD Bioscience |
| Perforin | FITC | pf344 | MabTech |
| T-bet | PerCP-Cy5.5 | 4B10 | eBioscience |
| CXCR5 | PE-Cy7 | MU5UBEE | eBioscience |
| CXCR5 | SB702 | MU5UBEE | eBioscience |
| CXCR5 | PE-eFluor610 | MU5UBEE | eBioscience |
| TIGIT | PE-Cy7 | MBSA43 | eBioscience |
| CD8 | BV650 | RPA-T8 | Biolegend |
| CD8 | BV785 | RPA-T8 | Biolegend |
| ICOS | APC-Cy7 | C398.4A | Biolegend |
| CD14 | BV510 | M5E2 | Biolegend |
| CD16 | BV510 | 3G8 | Biolegend |
| CD20 | BV510 | 2H7 | Biolegend |
| CD69 | BV605 | FN50 | Biolegend |
| PD1 | BV711 | EH12.2H7 | Biolegend |
| PD1 | BV785 | EH12.2H7 | Biolegend |
| PD1 | BV421 | EH12.2H7 | Biolegend |
| CX3CR1 | PE | K0124E1 | Biolegend |
| CD4 | APC-Cy7 | OKT4 | Biolegend |
| NKG2A | PE-Cy7 | Z199 | Beckman Coulter |
| CD28 | ECD | CD28.2 | Beckman Coulter |
| CD103 | APC-Cy7 | 2G5 | Beckman Coulter |

**Supplementary Figure 1:**
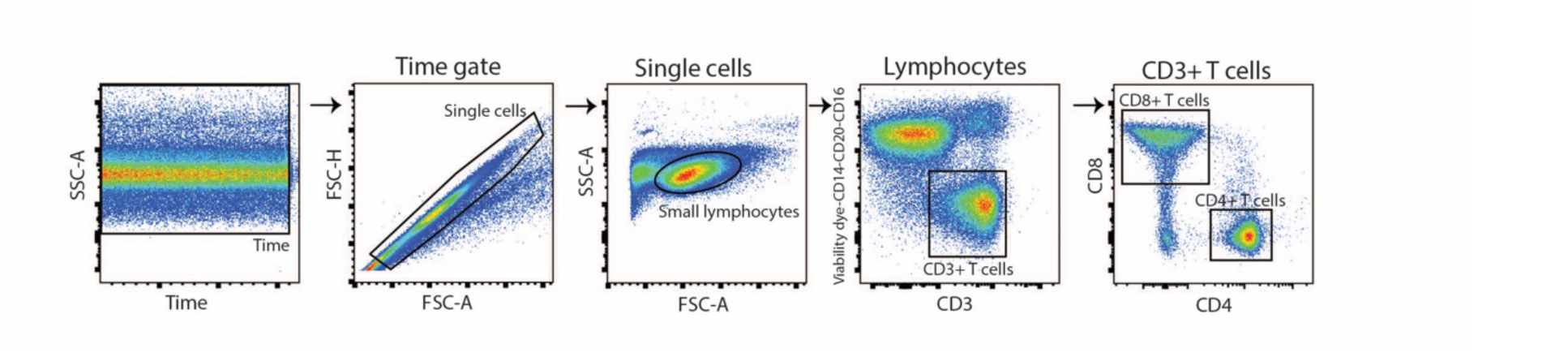
Gating strategy used for flow cytometric analyses. Representative example of a RM is shown. First, to ensure that only live single cells were analyzed from PBMCs, LNMCs or immune cells from the rectal mucosa, forward scatter height (FSC-H)-versus-forward scatter area (FSC-A) and side scatter area (SSC-A)-versus-FSC-A plots were used to exclude doublets and focus on singlet small lymphocytes. Dead cells were excluded by gating on cells negative for the viability marker Aqua Blue. Monocytes, B and NK cells were excluded via the CD14/20/16 dump gate. CD4^+^ and CD8^+^ T lymphocytes were gated within CD3^+^ cells. Following gates are shown in each corresponding figure.

**Supplementary Figure 2:**
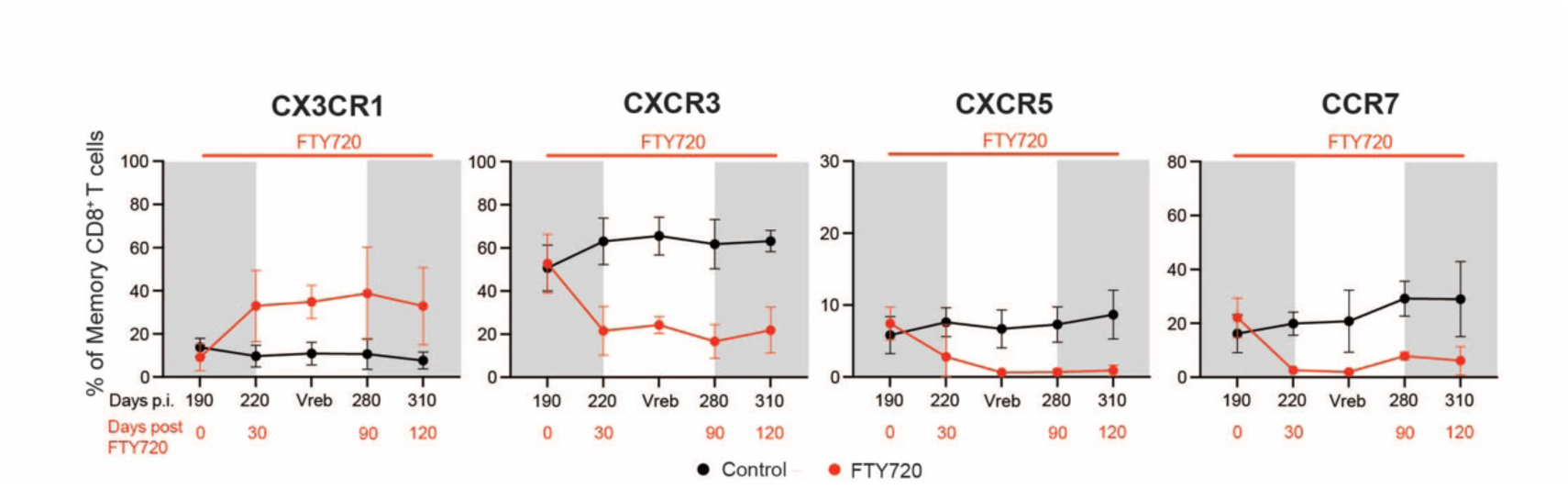
Phenotype of memory CD8+ T cells in blood during FTY720 administration. Expression of chemokine receptors CX3CR1, CXCR3, CXCR5 and CCR7 on circulating memory CD8+ T cells of control or FTY720-treated RMs at different time points. Grey rectangles indicate time points under ART. N=7 for each group for every time point analyzed except at day 220 p.i. where FTY720-treated group n=5. Frozen PBMCs where used to assess T cell phenotype except for CCR7 staining that was done on fresh samples. Data are presented as the mean ± SD.

**Supplementary Figure 3:**
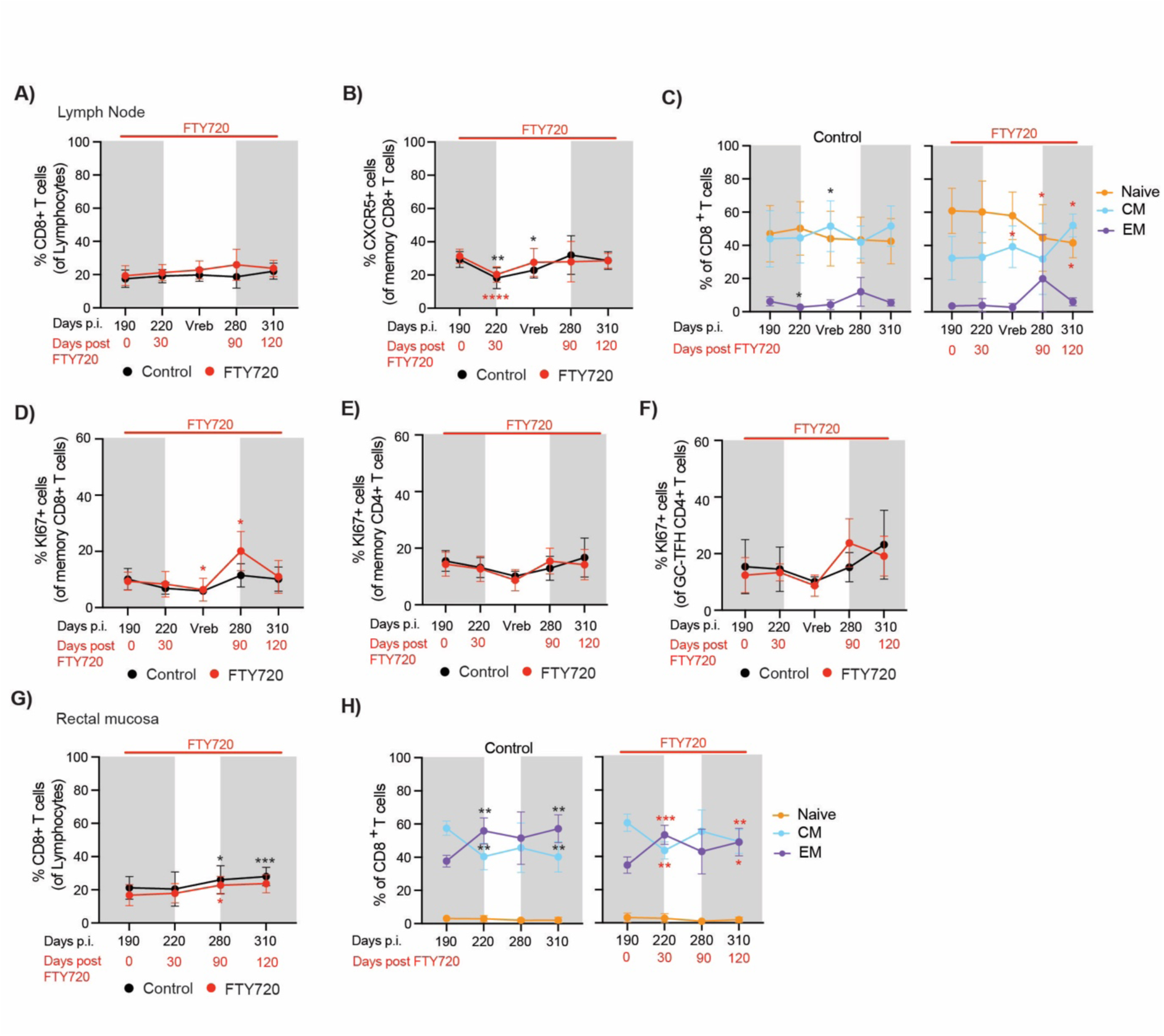
Proportion of CD8+ T cells and memory phenotype in lymph node and rectal tissue during FTY720 administration. A) Proportion of total CD8+ T cells, B) Proportion of CXCR5+ memory CD8+ T cells; and C) proportion of memory subsets (Naïve, CM -central memory- and EM -effector memory-) within CD8+ T cells in LN of control and FTY720-treated RMs over time. Proportion of KI67+ cells within D) Memory CD8+ T cells, E) Memory CD4+ T cells and F) GC-TFH CD4+ T cells in LN of control and FTY-720-treated RMs over time (n=7 for each group at every time point except for day 280 and 310 p.i. where n=6 for FTY720-treated RMs). G) Proportion of CD8+ T cells and H) proportion of memory subsets (Naïve, CM and EM) within CD8+ T cells in RB of control and FTY720-treated RMs over time (n=7 for each group at every time point). Fresh cells from LN and RB were used for flow cytometric analysis. Grey rectangles indicate time points under ART. Data are presented as the mean ± SD. Statistical differences were assessed by applying a 2 way-anova or mixed-effects model (if there were missing values) with Dunnett’s for comparisons between time points within each arm (every time point vs day 190 p.i.; black p values correspond to statistical analysis within the control arm, and red p values within FTY720 arm). Data are presented as the mean ± SD. *P≤0.05, **P≤0.01, ***P≤0.001, ****P≤0.0001.

**Supplementary Figure 4:**
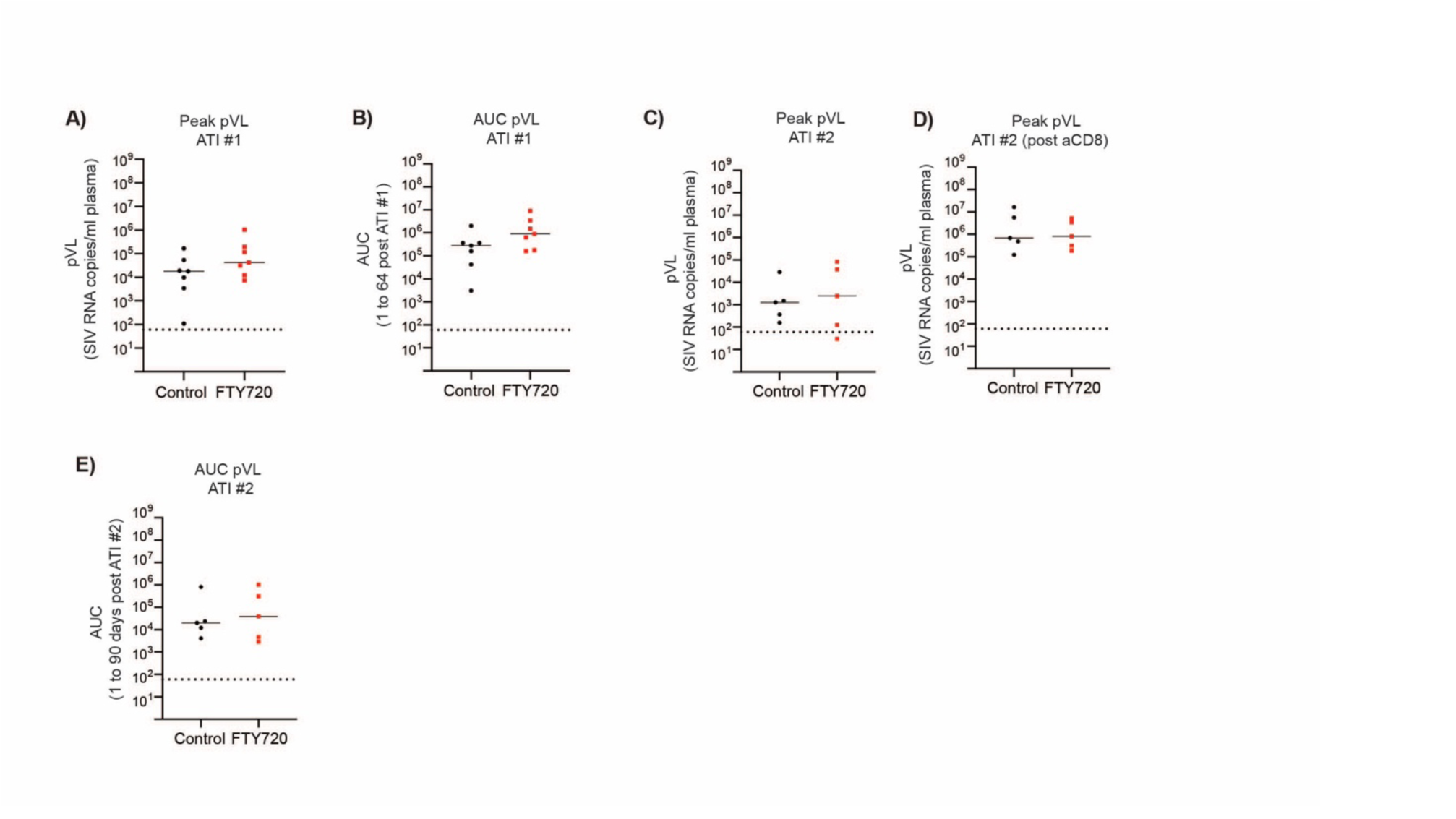
FTY720 does not impact peak viral load levels after first or second antiretroviral treatment interruption. Plasma SIVmac239M RNA levels expressed as copies/ml (limit of detection = 60 copies/ml, indicated with a dashed line) are shown for each individual animal from control (black dots) or RMs treated with FTY720 from 190 to 310 days p.i. (red dots). A) Peak plasma viral load during first ATI. B) Area Under the Curve at ATI#1. C) Peak plasma viral load during ATI#2, before CD8 depletion. D) Peak plasma viral load during ATI#2 after CD8 depletion. E) Area under the curve at ATI#2. Lines correspond to median values. Statistical differences were assessed with Mann-Whitney.

**Supplementary Figure 5:**
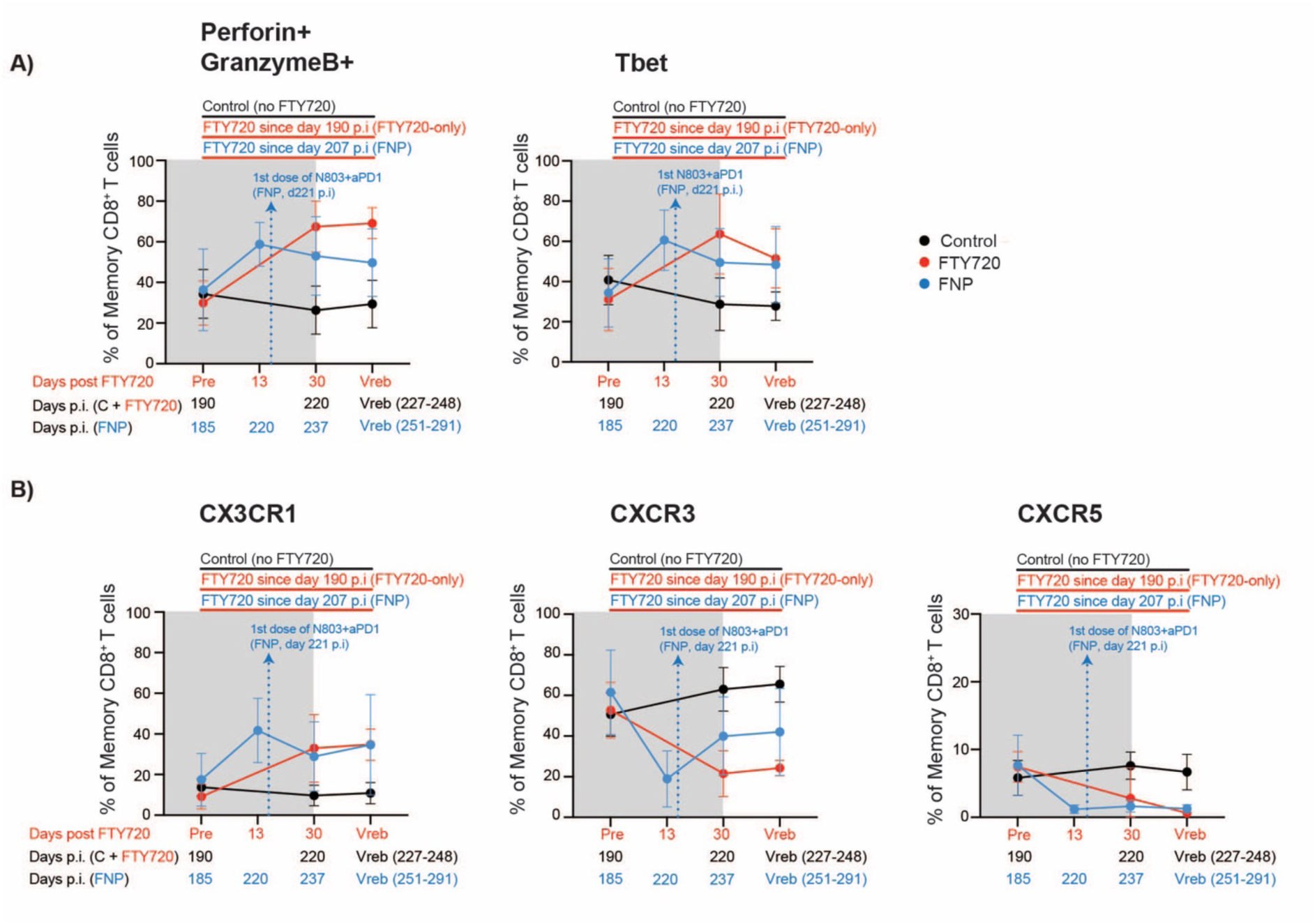
Phenotype of memory CD8+ T cells in blood FNP-treated RM. Expression of A) cytotoxic granules perforin and granzyme B and transcription factor Tbet; and B) chemokine receptors on circulating memory CD8+ T cells of control, FTY720-only or FNP-treated RMs at different time points equivalents between arms. Grey rectangles indicate time points under ART. N=7 for each group for every time point analyzed except at day 220 p.i. where FTY720-treated group N=5. Frozen PBMCs where used to assess T cell phenotype. Data are presented as the mean ± SD.

**Supplementary Figure 6:**
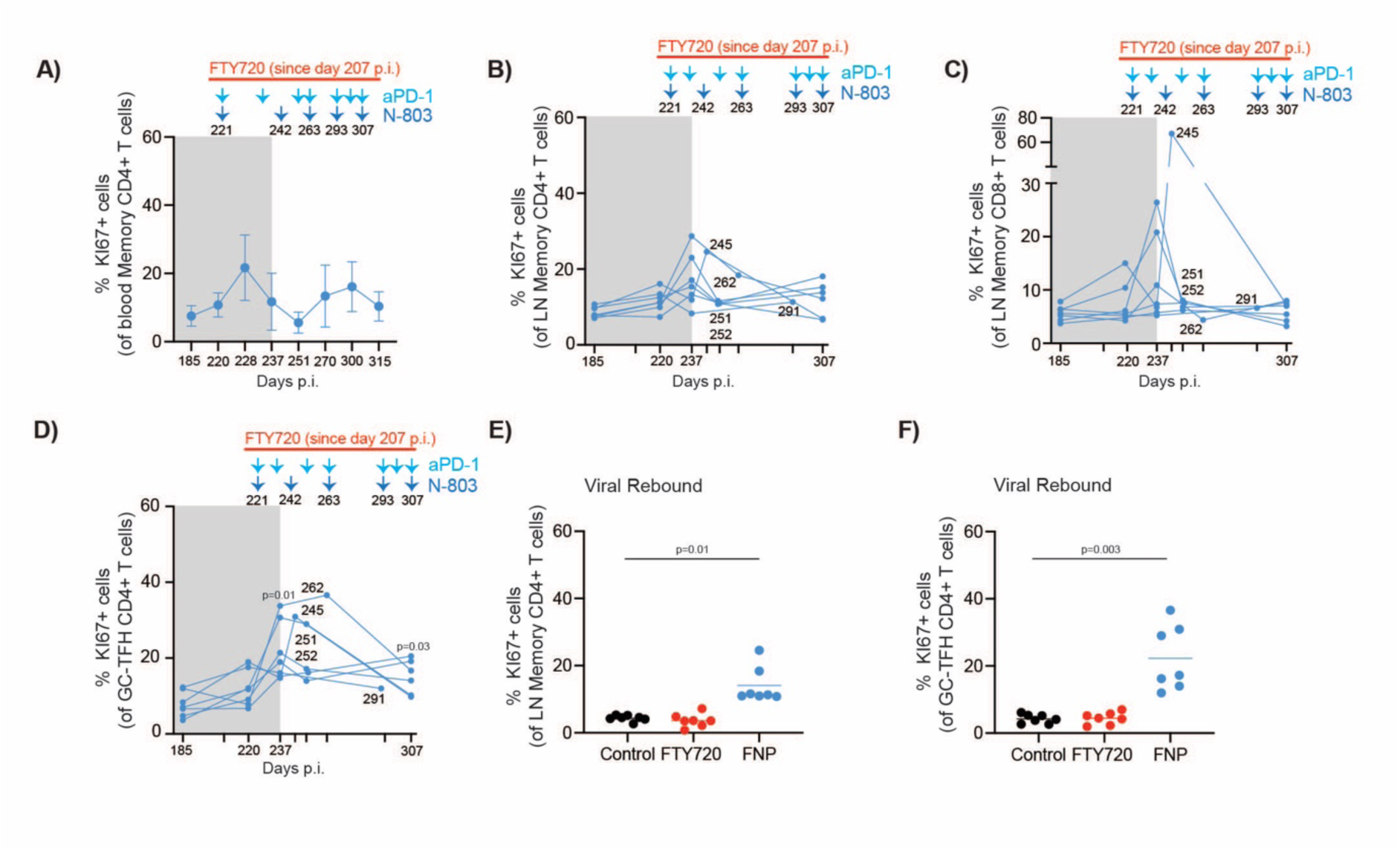
N-803 and aPD1 administration increased LN CD4+ T cell activation. A) Summary plot showing the changes in KI67 (surrogate for proliferation) in blood CD4+ T cells over time. Arrows indicate aPD1 and/or N-803 administration. Numbers below the arrows are days p.i. Mean and SD values are shown. B), C) and D) summary plots show the proportion of KI67+ cells within LN B) CD4+ T cells, C) CD8+ T cells and D) Germinal center follicular helper (GC-TFH) CD4+ T cells. Arrows indicate aPD1 and/or N-803 administration. Numbers below the arrows are days p.i. Grey rectangles indicate ART. E) and F) show the proportion of KI67+ cells within bulk LN CD4+ T cells (E) and GC-TFH CD4+ T cells (F) for control, FTY720-only or FNP-treated RM during ATI at viral rebound. N=7 for each group at each time point except for day 307 p.i for the FNP group where n=6. Lines correspond to median values. Cryopreserved mononuclear cells were used to assess T cell phenotype. Statistical differences in A), B), C and D) were calculated using one-way anova test with Dunnet’s multiple comparisons test vs pre-treatment time point (day 220 p.i for A) and day 185 for the rest). In B), C) and D) time points included in the statistical analysis were 185, 220, 237 and 307 p.i.. Satistical differences in E) and F) were assessed using Kruskal-wallis test with Dunn’s post comparisons versus control group.

